# Dysregulated Microglial Synaptic Engulfment in Diffuse Midline Glioma

**DOI:** 10.64898/2025.12.22.696064

**Authors:** Rebecca Mancusi, Eva Tatlock, Kiarash Shamardani, Lehi Acosta-Alvarez, Richard Drexler, Vrunda Trivedi, Avishai Gavish, Kouta Niizuma, Neeraj Soni, Pamelyn Woo, Sara Mulinyawe, Samin Jahan, Natalie Logan, Karen Malacon, Youkyeong Gloria Byun, Anna Geraghty, Tara Barron, Kathryn Taylor, Michelle Monje

**Author notes:** Corresponding author: Michelle Monje.

## Abstract

Diffuse midline glioma (DMG) is a near-universally lethal form of pediatric high-grade glioma, driven by neuronal activity-regulated paracrine signaling and synaptic integration of malignant cells into neural circuits. In turn, DMG increases neuronal excitability, augmenting neuron-to-glioma signaling. In the healthy brain, microglia, the resident immune cells of the central nervous system (CNS), regulate neuronal excitability and synaptic connectivity. However, the role of microglia in promoting tumor-associated hyperexcitable neural networks in glioma remains unknown. Here, we investigate the activity-regulated engulfment of neuronal synapses by microglia in both healthy and glioma-bearing mice, and further explore how glioma cells alter microglia-mediated circuit refinement, contributing to pathogenic neuronal hyperexcitability. Microglia-mediated circuit refinement in the glioma microenvironment was characterized through synaptic engulfment analysis of both excitatory and inhibitory synapses by microglia in healthy mice and patient-derived DMG xenograft models, paired with optogenetic stimulation in the neocortex. We found that glutamatergic neuronal activity in the healthy brain increased excitatory synaptic engulfment by microglia in a previously unappreciated negative feedback mechanism that may guard against hyperexcitability. In contrast, this activity-regulated increase in excitatory synaptic engulfment was abrogated in DMG-infiltrated brains. Instead, inhibitory synaptic engulfment was significantly increased in DMG in response to glutamatergic neuronal activity. Together, these dysregulated synaptic engulfment mechanisms may create imbalance in the excitatory to inhibitory (E:I) synapse ratio predicted to increase neuronal excitability. Complementary single-nuclei sequencing studies revealed concordant tumor-specific, activity-regulated changes in microglia-neuron signaling showing reduced expression of excitatory synaptic refinement gene programs in microglia, potentially mediating the aberrant synaptic engulfment observed in DMG. These findings reveal novel cancer-neuron-immune interactions in DMG and provide an opportunity to potentially modulate tumor-associated neuronal hyperexcitability by targeting aberrant microglial synaptic engulfment.

**HIGHLIGHTS:**

- Excitatory synaptic engulfment by microglia increases in a glutamatergic neuronal activity-dependent manner in the healthy brain.
- In the glioma-infiltrated brain, this feedback regulatory mechanism is inverted, with loss of activity-regulated excitatory synaptic engulfment and aberrant increased inhibitory synaptic engulfment.
- Transcriptomic changes in glioma-associated microglia contribute to differential excitatory and inhibitory synaptic engulfment, with reduced excitatory synaptic refinement gene programs in tumor-associated microglia.
- Activity-regulated soluble factors drive this imbalanced excitatory-inhibitory postsynaptic engulfment by glioma-associated microglia.

## INTRODUCTION

Pediatric high-grade gliomas (pHGGs) are the leading cause of cancer-related deaths in children^1^. These brain cancers, including H3K27M-altered diffuse midline gliomas (DMG), exhibit unique spatiotemporal patterns^2^, with pontine (also called diffuse intrinsic pontine glioma, DIPG) and thalamic diffuse midline gliomas typically presenting in mid-childhood and multiple subtypes of high-grade cerebral hemispheric gliomas occurring more commonly in adolescence and young adulthood^3,4^. They arise during distinct periods of postnatal neurodevelopment, suggesting neurodevelopmental origins^5, 6^. DMGs, in particular, originate from early oligodendroglial precursor cells (OPCs)^5–7^, that are known to proliferate in response to neuronal activity^8–9^. In DMG and other gliomas, neuronal activity similarly drives tumor growth^10–11^ and initiation^12^ through activity-regulated secreted factors^10, 13, 14^ and bona fide neuron-to-glioma synapses^14–18^, highlighting the crucial significance of the brain microenvironment in DMG pathobiology. A prominent feature of dysregulated neuron-glioma interactions in the glioma microenvironment is tumor-associated neuronal hyperexcitability, driven by the secretion of glutamate^19, 20^ and synaptogenic factors^17, 18, 21–24^ that perpetuates a feed-forward cycle of glioma-growth-promoting neuronal activity.

While the critical role of neuronal activity in driving glioma proliferation has been increasingly recognized, the involvement of glial cells in the tumor microenvironment and their contribution to pathogenic neuron-to-glioma networks remains understudied. Microglia, the brain’s resident immune cells, are essential for neural circuit formation and function. In the healthy brain, microglia assume a variety of cell states^25–28^ to dynamically interact with synapses^29, 30^, modulate neuronal excitability^31–33^ and regulate synapse refinement to promote proper circuit connectivity during development^34–45^ and throughout life^46–51^. Microglia comprise a significant portion of the brain tumor microenvironment^52–55^, with pediatric glioma-associated microglia exhibiting unique transcriptional states compared to their adult counterparts^56^. These glioma-associated microglia contribute to the glioma microenvironment by secreting factors that promote tumor growth, including IL-6^57, 58^, TGF-β^56, 59–61^, and others^62–67^, and can potentially be therapeutically targeted via CSF1R signaling to reduce tumor growth^68, 69^. Notably, these glioma-associated microglia do not conform to traditional pro- and anti-inflammatory phenotypes^70, 71^, suggesting that they adopt a distinct functional state that supports glioma progression. Recent transcriptomic analyses of pHGG-associated microglia have further underscored their role in tumor growth, with these cells expressing substantially less pro-inflammatory cytokines like CCL3/4^56^ compared to microglia in adult glioblastoma (GBM). While these studies highlight the importance of microglia in pHGG, possible contributions to pathogenic neuron-to-glioma networks remain largely unexplored.

The balance between excitatory and inhibitory synapses is crucial for maintaining neural circuit homeostasis and function and preventing hyperexcitability^72–77^. Recent studies have highlighted hyperexcitable neural circuits in both adult^19, 21–24^ and pediatric gliomas^15, 24^, revealing an increase in excitatory synapses ^21, 23^, reduction in inhibitory synapses^78^, alongside elevated neuronal firing rates and amplitudes within these remodeled circuits^15, 19, 22, 78^. Microglia play a key role in the regulation of excitatory and inhibitory synaptic engulfment during postnatal development^34–44^, throughout life^48, 49^, and in disease ^47, 79–81^, while dysregulated synaptic engulfment can contribute to neurological and psychiatric dysfunction^31, 79, 80, 82–89^. Microglia utilize a variety of molecular machinery to mediate excitatory synaptic engulfment, including the complement pathway^34, 37^ and P2RY12 cascade^31^, which can become dysregulated to drive pathological conditions^31, 46, 79, 86, 87^. Microglia additionally facilitate inhibitory synaptic engulfment, for example through GABA ^38^ and progranulin signaling^83^, which similarly becomes impaired in pathological contexts^84, 90^. However, the contribution of putatively altered synaptic engulfment by microglia to glioma-induced neural circuit modifications has yet to be fully explored. Here, we demonstrate that microglia modulate excitatory and inhibitory synaptic engulfment in a neuronal activity-driven manner in both healthy and glioma-bearing mice.

## RESULTS

### Synaptic engulfment by microglia is increased in diffuse midline glioma

In order to understand how microglia may pathogenically sculpt neural circuits and contribute to neuronal hyperexcitability, we utilized a patient-derived orthotopic xenograft model of DMG (‘SU-DIPG-VI’) or a vehicle control (‘Nontumor control’, buffered saline) in the M2 frontal cortex region of 4-week-old NOD-SCID-IL2-gamma chain-deficient (NSG) mice (**Fig. 1A**). Notably, while DMGs initially arise in midline structures, they rapidly infiltrate neural circuits throughout the brain, including the densely innervated cortex^55, 91^.

**Figure 1:**
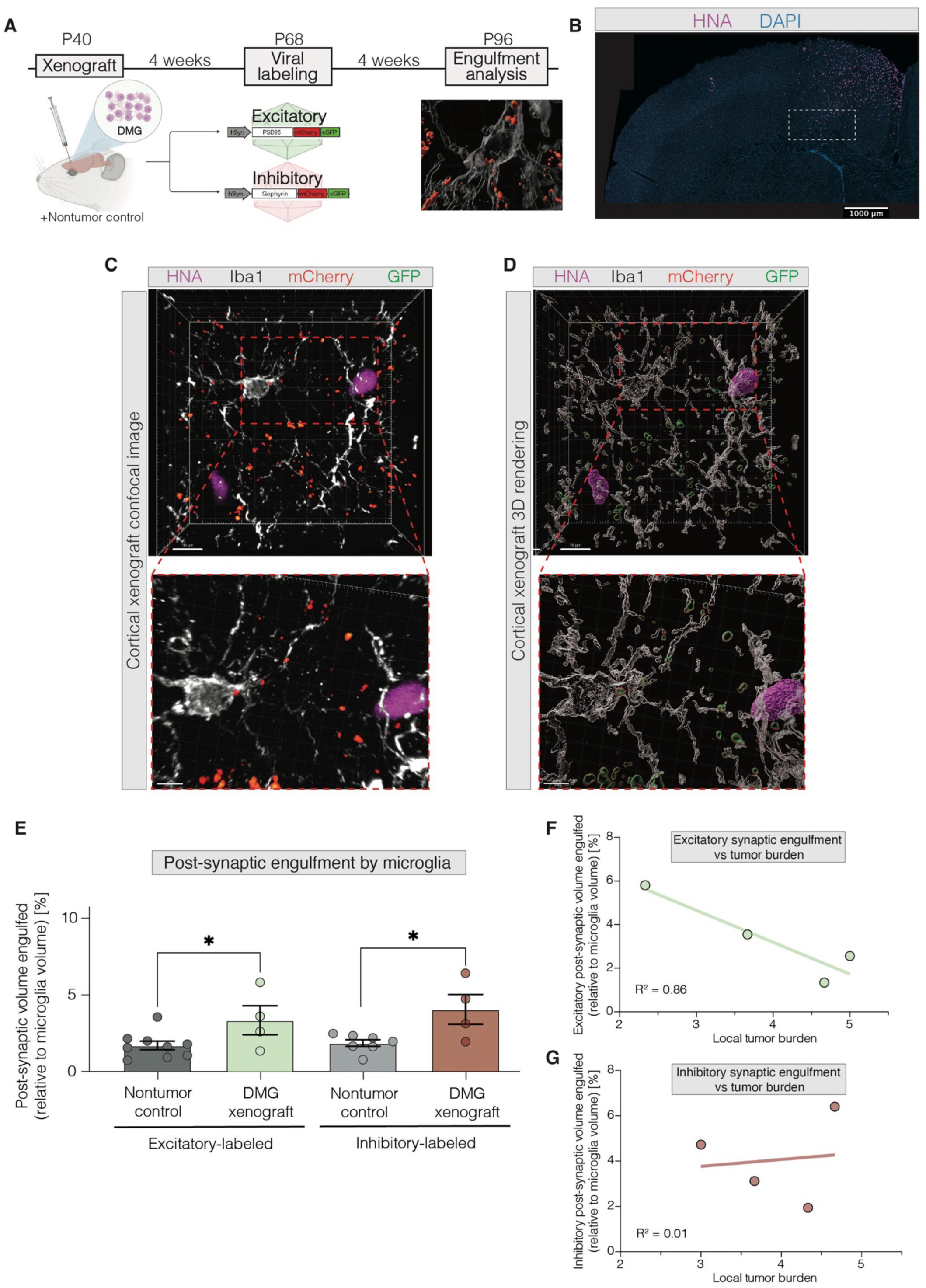
Synaptic engulfment by microglia is increased in diffuse midline glioma. (A) Xenograft and synaptic labeling paradigm in NOD scid gamma (NSG) mice which underwent cortical xenograft with DMG, diffuse intrinsic pontine glioma (patient-derived cell line “SU-DIPG-VI”), or saline injections (“Nontumor control”), followed by viral injections with hSyn-PSD95-mCherry-eGFP (“Excitatory”) AAV9 or hSyn-gephyrin-mCherry-eGFP (“Inhibitory”) AAV9 in mouse premotor frontal cortex (M2). Synaptic engulfment analysis was then performed. P, postnatal day; hSyn, human synapsin promoter. (B) Representative histological images of glioma (SU-DIPG-VI) xenografted into mouse cortex. DAPI (4’,6-diamidino-2-phenylindole, blue) marks nuclei, HNA (human nuclear antigen, violet) marks glioma cells. Subsequent analyses performed in diffusely infiltrating edge of DMG xenograft, outlined in white. Scale bar, 1000 um. (C) Representative histological and corresponding 3D rendered images (D) of HNA+ glioma cells (SU-DIPG-VI, violet), Iba1+ microglia (white), mCherry+ gephyrin+ synapses (red), and GFP+ gephyrin+ synapses (green) in xenografted deep-layer premotor cortex. Scale bar, 10 um. (E) Volumetric engulfment analysis of microglial engulfment of mCherry+ PSD95+ synapses (“Excitatory-labeled”) and mCherry+ gephyrin+ synapses (“Inhibitory-labeled”), represented as the % volume of mCherry+ synapse contained in Iba1+ microglia (as in representative image in B) in either saline injected (“Nontumor control”, grey) or glioma xenografted (“DMG xenograft”, SU-DIPG-VI, green or red) mice. (F-G) Correlative analysis of microglial engulfment of excitatory- (F, green) or inhibitory-labeled (G, red) synapses (as in E), compared to average number of tumor cells in analyzed regions (“Local Tumor Burden”). Each individual point represents the average volumetric quantification of synaptic engulfment (y-axis, as in E) and corresponding number of tumor cells (x-axis) analyzed in one mouse. Each individual point represents the average volumetric quantification from images analyzed in one mouse (E-G). Data are mean ± s.e.m. One-way ANOVA with Tukey’s post hoc analysis (E); *P<0.05. R^2^ values determined using linear regression model.

After 4 weeks of tumor engraftment, excitatory and inhibitory postsynaptic structures were fluorescently labeled using previously established viral vectors to mark excitatory or inhibitory synapses^92^. DMG-bearing and nontumor control mice were injected with either an “excitatory synapse” viral vector, using the hSyn promoter to drive expression of PSD95–mCherry–eGFP in excitatory synapses (AAV9-CW3SL::hSyn-PSD95-mCherry-eGFP), or an “inhibitory synapse” viral vector, using hSyn to drive expression of gephyrin–mCherry–eGFP in inhibitory synapses (AAV9-CW3SL::hSyn-gephyrin-mCherry-eGFP)^92^. Both mCherry and eGFP maintain fluorescence under neutral pH conditions, whereas only mCherry signal is preserved under acidic pH conditions, such as those found in phagocytic lysosomes. This vector design enables robust monitoring of glial engulfment of synapses *in vivo*^92^.

Mice were perfused 8 weeks after xenografting and 4 weeks after viral expression of synaptic markers targeting neurons. High-resolution confocal microscopy imaging was conducted at the diffusely infiltrating edge of the DMG xenograft (**Fig. 1B**). Microglial engulfment of mCherry-labeled synapses was observed within the DMG tumor microenvironment. Notably, microglia actively engulfing synapses were visualized wrapping their processes around adjacent DMG cells (**Fig. 1C-D**). 3D rendering was then performed to compute the volumes of all labeled objects (**Fig. 1D, Supp. Video 1**).

Following established quantification methods for microglial synaptic engulfment^34, 37^, we quantified the microglial engulfment of excitatory and inhibitory postsynaptic structures (see STAR methods). We found that microglia in DMG-xenografted mice engulfed significantly more excitatory postsynaptic structures compared to microglia in nontumor controls (**Fig. 1E**). Similarly, microglia in DMG-xenografted mice also engulfed significantly more inhibitory postsynaptic structures than those in the nontumor control mice (**Fig. 1E**). These findings suggest that microglia in the context of DMG generally adopt an enhanced ability to engulf synapses.

We further investigated whether excitatory and inhibitory postsynaptic engulfment by microglia varied in a tumor-burden-dependent manner. Our analysis revealed that microglia engulfed significantly fewer mCherry^+^ excitatory postsynaptic structures as the local tumor burden increased (**Fig. 1F**). In contrast, microglial engulfment of inhibitory synapses remained unchanged in response to local tumor burden (**Fig. 1G**). This discrepancy in tumor-burden-dependent synaptic engulfment suggests that DMG-associated microglia may differentially modulate excitatory and inhibitory synapses, positioning glioma-associated microglia as possible mediators of E:I imbalance in DMG-affected neural circuits.

### Activity-dependent excitatory synaptic engulfment in healthy cortex

We next investigated whether neuronal activity regulates excitatory and inhibitory synaptic engulfment by microglia in the healthy and glioma-infiltrated cortex. To address this, we xenografted patient-derived DMG (‘SU-DIPG-VI’) cells or vehicle control (‘Nontumor control’) into the M2 cortical region of 4-week-old Thy1::ChR2 NSG mice, which express the excitatory, blue-light-gated opsin channelrhodopsin-2 in deep-layer cortical projection neurons^93^ (**Fig. 2A**). Following 4 weeks engraftment, mice were injected with the aforementioned AAV9 vectors^92^ to label either postsynaptic excitatory or inhibitory synapses. An optical ferrule was placed in superficial cortical layers of the M2 frontal cortex in DMG-xenografted and nontumor control mice. After 4 weeks of viral expression to mark excitatory or inhibitory synapses, optogenetic stimulation of glutamatergic cortical projection neurons (one 30-min session, 20 Hz blue-light stimulation, with 30-s on/90-s off cycles)^8, 10, 14, 94^ was performed. DMG-xenografted and nontumor groups of littermate controls that did not express ChR2 were identically manipulated to control for light exposure and optical ferrule surgical manipulation. Mice were sacrificed 24 hours after optogenetic stimulation.

**Figure 2:**
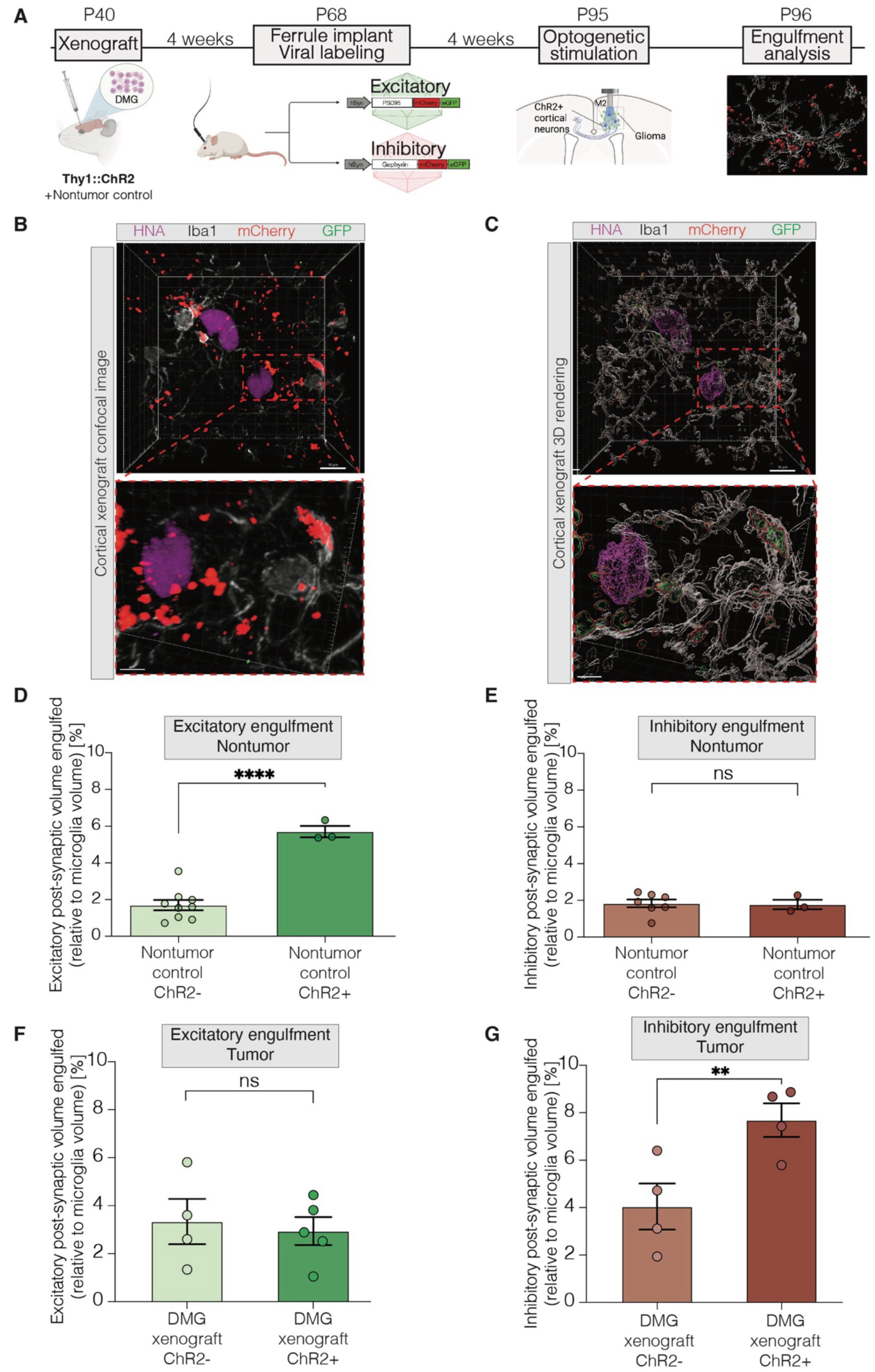
Neuronal activity-dependent engulfment of excitatory synapses in the healthy cortex is abrogated in glioma. (A) Xenograft, synaptic labeling, and optogenetics paradigm in NOD scid gamma (NSG) mice which underwent cortical xenograft with DMG, diffuse intrinsic pontine glioma (patient-derived cell line “SU-DIPG-VI”), or saline injections (“Nontumor control”), followed by viral injections with hSyn-PSD95-mCherry-eGFP (“Excitatory”) AAV9 or hSyn-gephyrin-mCherry-eGFP (“Inhibitory”) AAV9 and optogenetic ferrule implantation in mouse premotor frontal cortex (M2). NSG mice used were either positive or negative for Thy1::ChR2 transgene. One acute session of optogenetic stimulation was performed within DMG-xenografted or saline-injected M2 cortical regions, according to previously-established methods, on Thy1::ChR2 +/− cohorts. Synaptic engulfment analysis was then performed. P, postnatal day; hSyn, human synapsin promoter. (B) Representative histological and corresponding 3D rendered images (C) of HNA+ glioma cells (SU-DIPG-VI, violet), Iba1+ microglia (white), mCherry+ gephyrin+ synapses (red), and GFP+ gephyrin+ synapses (green) in xenografted deep-layer premotor cortex of Thy1::ChR2 negative (“ChR2-”) mice. Scale bar, 10 um. (D) Volumetric engulfment analysis of microglial engulfment of mCherry+ PSD95+ synapses (“Excitatory”), represented as the % volume of mCherry+ synapse contained in Iba1+ microglia in saline injected (“Nontumor control”) mice without Thy1::ChR2 expression (“ChR2-”) or with Thy1::ChR2 expression (“ChR2+”). (E) Volumetric engulfment analysis of microglial engulfment of mCherry+ gephyrin+ synapses (“Inhibitory”), represented as the % volume of mCherry+ synapse contained in Iba1+ microglia in saline injected (“Nontumor control”) mice without Thy1::ChR2 expression (“ChR2-”) or with Thy1::ChR2 expression (“ChR2+”). (F) Volumetric engulfment analysis of microglial engulfment of mCherry+ PSD95+ synapses (“Excitatory”), represented as the % volume of mCherry+ synapse contained in Iba1+ microglia in glioma xenografted (“DMG xenograft”, SU-DIPG-VI) mice without Thy1::ChR2 expression (“ChR2-”) or with Thy1::ChR2 expression (“ChR2+”). (G) Volumetric engulfment analysis of microglial engulfment of mCherry+ gephyrin+ synapses (“Inhibitory”), represented as the % volume of mCherry+ synapse contained in Iba1+ microglia in glioma xenografted (“DMG xenograft”, SU-DIPG-VI) mice without Thy1::ChR2 expression (“ChR2-”) or with Thy1::ChR2 expression (“ChR2+”). Each individual point represents the average volumetric quantification from images analyzed in one mouse (B-C, F-G). Data are mean ± s.e.m. One-way ANOVA with Tukey’s post hoc analysis (B-C, F-G); **P < 0.01, ****P < 0.0001; NS, not significant.

In order to better understand activity-regulated patterns of synaptic engulfment, we performed similar high-resolution confocal microscopy analyses of microglial synaptic engulfment (**Fig. 2B,C**). In non-tumor control mice, we found that microglia in deep layer M2 frontal cortex engulf significantly more excitatory synapses in response to cortical neuronal activity induced by acute optogenetic stimulation of deep layer cortical projection neurons (**Fig. 2D**), highlighting a previously under-recognized negative feedback mechanism of microglial excitatory synaptic engulfment. Microglial synaptic engulfment of inhibitory synapses remained unchanged by acute optogenetic stimulation of glutamatergic neurons (**Fig. 2E**).

### Dysregulation of activity-dependent excitatory synaptic engulfment in glioma-infiltrated cortex

In stark contrast, in the presence of DMG cells infiltrating frontal cortex, microglia do not increase excitatory synaptic engulfment in response to local glutamatergic neuronal activity (**Fig. 2F**). Instead, microglia engulf more inhibitory synapses in response to acute optogenetic stimulation of glutamatergic neurons in DMG-bearing mice (**Fig. 2G**).

Taken together, these findings suggest that in the healthy brain, glutamatergic neuronal activity induces a feedback mechanism of excitatory - but not inhibitory - synaptic engulfment that may maintain homeostatic circuit excitability. In the glioma-infiltrated cortex, DMG-associated microglia become dysregulated and instead engulf inhibitory rather than excitatory synapses in response to glutamatergic neuronal activity.

### Neuronal activity alters microglial transcriptomic states and circuit refinement-related pathways in glioma

In order to enable deeper molecular characterization in a fully immunocompetent model, we utilized a murine DMG allograft model that leverages H3K27M mosaic analysis with double recombination, “MADR” ^95^. We first sought to characterize the electrophysiological properties of MADR DMG. Field potentials recorded from mice with CA1 hippocampal murine DMG allografts (**Fig. 3A-B**), displayed enhanced field excitatory postsynaptic potentials across a series of current injections compared to the nontumor controls, in line with previous electrophysiological characterizations of neuronal hyperexcitability in DMG ^15^ and adult GBM models^19, 20^. As observed in DMG models^10, 13, 14, 16^, murine glioma MADR cells increase proliferation in response to conditioned media from optogenetically-stimulated Thy1::ChR2+ ex vivo brain slices-termed active conditioned media, or “ACM”-(**Supp. Fig. 1A**) compared to conditioned media from nonstimulated Thy1::ChR2-ex vivo brain slices-termed conditioned media or “CM“-. As occurs in patients with DMG^55^, the MADR model of DMG diffusely infiltrates cortex – evident histologically (**Supp. Fig. 1B-C**) and via magnetic resonance imaging (MRI) (**Supp. Fig. 1D**).

**Figure 3:**
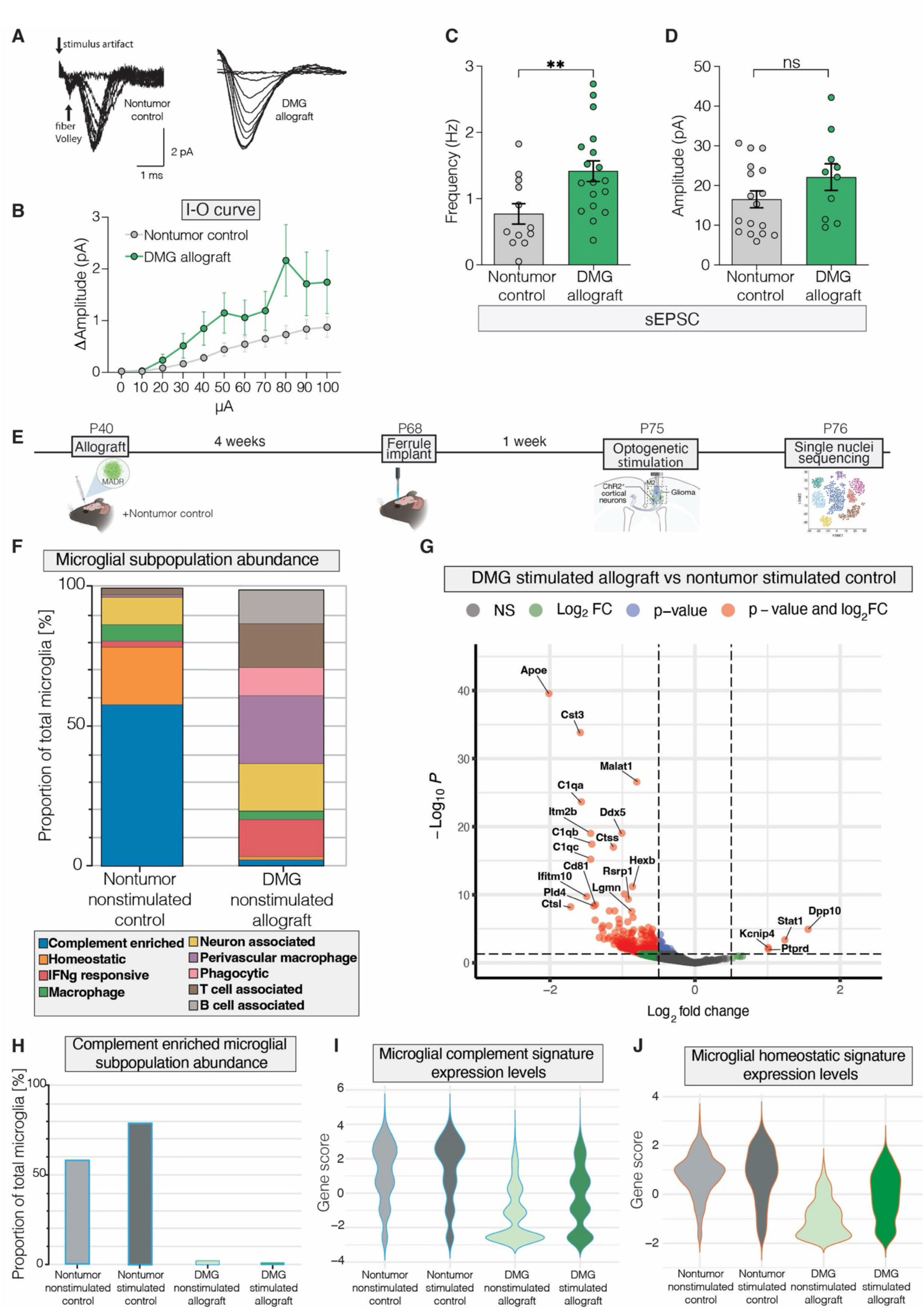
Neuronal activity alters microglial populations, modulating circuit refinement-related pathways in glioma. (A) Repetitive superimposed traces of field potentials recorded from mice with CA1 hippocampal murine glioma MADR allografts (“DMG allograft”) or from control saline-injected mice (“Nontumor control”). (B) Input-output (I-O) curve was performed in the CA1 region of hippocampal slices. Field excitatory postsynaptic potentials (fEPSPs) were recorded from mice with CA1 hippocampal murine glioma MADR allografts (“DMG allograft”) or from control saline-injected mice (“Nontumor control”). Changes in amplitude were measured relative to baseline following a series of current injections ranging from 0 to 140 µA in steps of 10 µA. (C-D) Frequency and amplitude of spontaneous excitatory postsynaptic currents (sEPSCs) were measured in mice with CA1 hippocampal murine glioma MADR allografts (“DMG allograft”) or from control saline-injected mice (“Nontumor control”). (E) Allograft and optogenetics paradigm in CD-1::C57BL/6J (CD1::BL6) mice which underwent cortical allograft with previously-established MADR (“Mosaic analysis by dual recombinase-mediated cassette exchange”) murine glioma (H3.3K27M+ murine-derived cell line), or saline injections (“Nontumor control”), followed by optogenetic ferrule implantation in mouse premotor frontal cortex (M2). CD1::BL6 mice used were either positive or negative for Thy1::ChR2 transgene. One acute session of optogenetic stimulation was performed within MADR-allografted or saline-injected M2 cortical regions, according to previously-established methods, on Thy1::ChR2 +/− cohorts. Single nuclei sequencing analysis was then performed. P, postnatal day. (F) Microglial subpopulation abundance (labeled by color) was calculated as the % of total microglia (“Proportion of Total Microglia”, y-axis) in either nontumor nonstimulated control mice or glioma allografted nonstimulated mice (x-axis). (G) Scatterplot demonstrating gene expression changes in microglia from stimulated, glioma-allografted mice, as compared to stimulated, nontumor controls. The x-axis demonstrates log2 (fold change of stimulated glioma- vs nontumor-associated microglia) and the y axis demonstrates Log10P of the gene expression level. Points shown in red represent genes showing statistically significant changes. (H) Complement-enriched microglial subpopulation abundance was calculated as the % of total microglia (“Proportion of Total Microglia”, y-axis) across all experimental cohorts (x-axis). (I) Complement-associated gene score (y-axis) quantifies the relative expression levels of a complement-associated gene set in microglia across all experimental conditions (x-axis). (J) Homeostasis-associated gene score (y-axis) quantifies the relative expression levels of a homeostasis-associated gene set in microglia across all experimental conditions (x-axis). Data are presented as mean ± s.e.m. Unpaired t test with Welch’s correction (D-E); **P < 0.01; NS, not significant.

In line with this observed hyperexcitability in our DMG model, we identified an increase in the frequency of spontaneous excitatory postsynaptic currents (EPSCs) in tumor-associated neurons compared to the nontumor controls (**Fig. 3C**), with no significant difference in neuronal EPSC amplitude between groups (**Fig. 3D**); we did not observe a difference in either frequency or amplitude of inhibitory postsynaptic currents (IPSCs) in DMG-associated neurons (**Supp. Fig. 1E-F**).

To better understand the molecular changes associated with this DMG-associated pattern of activity-regulated microglial synaptic engulfment, we subsequently conducted a single-nuclei RNA sequencing (sn-RNAseq) analysis. We utilized the aforementioned MADR allograft model of murine DMG^95^ in the M2 region of the frontal cortex of 4-week-old Thy1::ChR2 CD1::BL6 mice, with vehicle (saline)-injected nontumor control littermates (**Fig. 3E**). After 4 weeks of engraftment, we placed an optical ferrule within the superficial layers of M2 cortex in DMG-allografted and nontumor controls. One week after ferrule placement, we optogenetically stimulated deep-layer glutamatergic cortical projection neurons in these Thy1::ChR2 mice as above. Mice were perfused 24 hours later, and deep cortical and striatal areas were then microdissected and processed for sn-RNAseq.

Various glial and neuronal cell types were identified in our computational analysis (**Supp. Fig. 2A**). We found multiple distinct microglial subpopulations (**Supp. Fig. 2B**), including complement-associated and neuron-associated microglia, which have been previously implicated in neuronal circuit refinement in development^25, 26^. While complement-associated microglia in our dataset are characterized by genes including *C1qa, C1qb, and C1qc* (**Supp. Table 1**), neuron-associated microglia express genes associated with microglia-neuron interactions including *Dpp10, Ptprd,* and *Nrg3* (**Supp. Table 1**). Quantification showed vastly different microglial subpopulations in DMG-allografted mice compared to nontumor, saline-injected controls (**Fig. 3F**). Strikingly, more homeostatic and complement-enriched microglia were found in nontumor saline-injected controls, whereas the phagocytic and neuron-associated microglial populations were enriched in the DMG-allografted mice (**Fig. 3F)**. We next examined glutamatergic activity-regulated gene expression changes in optogenetically stimulated, DMG-allografted mice compared to optogenetically stimulated, nontumor control mice (**Fig. 3G**). In response to glutamatergic cortical activity, DMG-associated microglia exhibit downregulation of complement pathway-associated genes and up-regulation of various genes associated with microglial-neuron interactions compared to nontumor control mice following optogenetic stimulation (**Fig. 3G)**.

Furthermore, complement-associated microglial subpopulations expand in response to glutamatergic neuronal activity in nontumor control mice (**Fig. 3H**). This indicates a dynamic activity-dependent expansion of complement-associated microglia populations in the healthy brain, while these populations are almost entirely lost in the DMG microenvironment (**Fig. 3H**). Complement-associated microglial gene expression levels across groups change in parallel with the observed differences in population abundance across groups (**Fig. 3I**). Additionally, homeostatic microglial gene expression becomes downregulated in the presence of DMG (**Fig. 3J)**. In line with these findings, the complement pathway has previously been shown to be critical for excitatory circuit refinement and synaptic remodeling in development^34, 37^ and disease^46, 79, 86^. The loss of complement pathway activation in DMG-associated microglia suggests that these tumor-associated microglia may lose the ability to sense and respond to the excitatory neuronal activity known to drive DMG progression^10, 13–15^.

### Glioma-infiltrated brain-derived and activity-regulated secreted factors affect microglial synaptic engulfment and activation state *in vitro*

Given the stark differences we observed in tumor-specific, activity-regulated microglial states *in vivo*, we next sought to understand the mechanisms mediating this effect and hypothesized that activity-regulated secreted factors may mediate this DMG-induced dysregulation of microglial synaptic engulfment. Media was collected from glioma-bearing or nontumor-bearing ex vivo brain slices that were either stimulated (Thy1::ChR2+, termed “active conditioned media” or “ACM”) or nonstimulated, spontaneously active (Thy1::ChR2-, termed “conditioned media” or “CM”). Tricultures of induced pluripotent stem cell (hiPSC)-derived microglial-like cells (iMGLs), hiPSC-derived GABAergic neurons and hiPSC-derived glutamatergic neurons were then exposed to conditioned medium from each of these four conditions (**Fig. 4A**). This experimental design also allows us to differentiate between engulfment of neuron-to-neuron synapses and potential engulfment of neuron-to-glioma synapses^15–17^ in the *in vivo* experiment above, as only neuron-neuron synaptic connections are present in this *in vitro* system.

**Figure 4:**
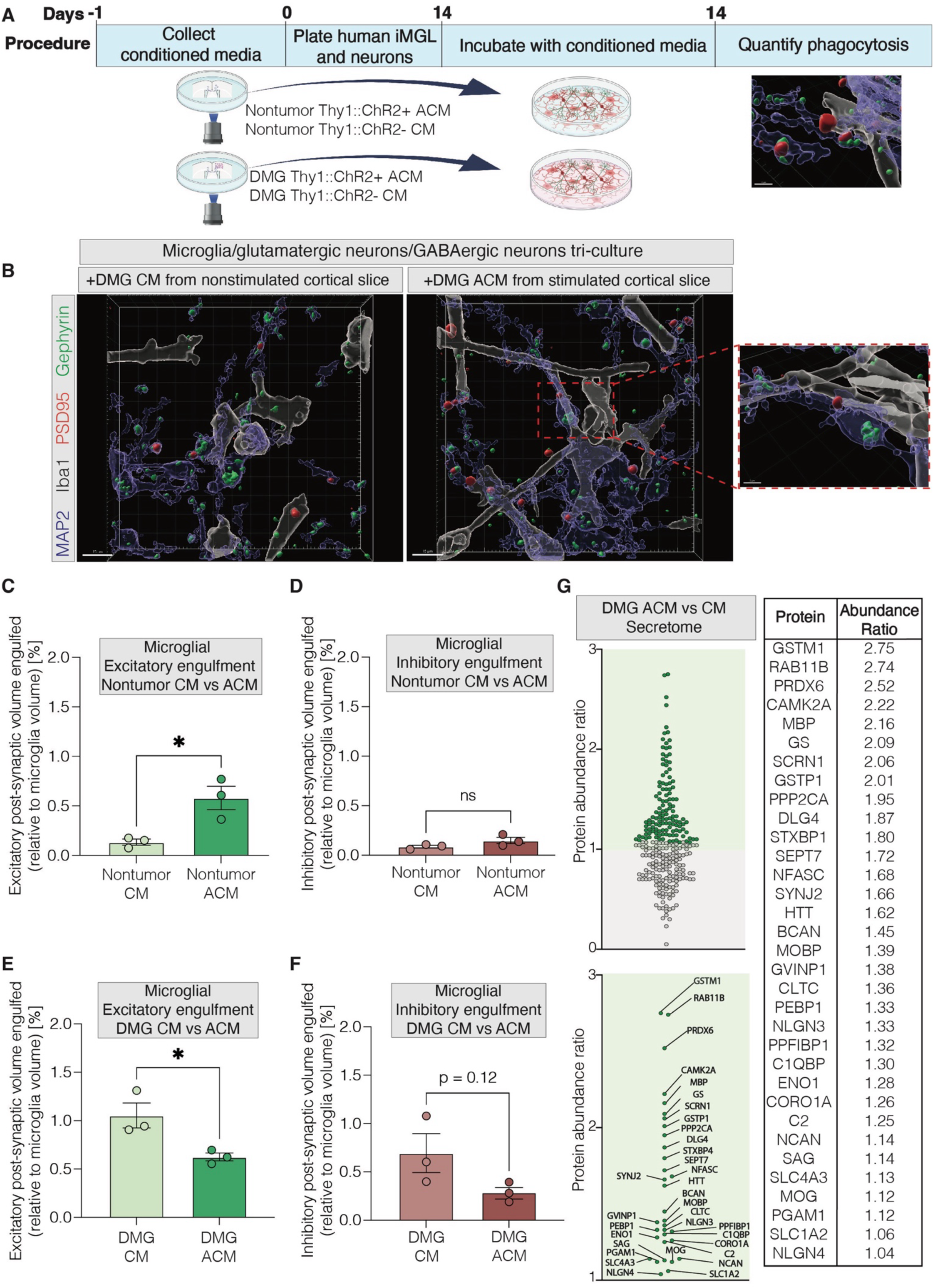
Glioma-infiltrated brain-derived and activity-regulated secreted factors affect microglial synaptic engulfment and activation state in vitro. (A) Paradigm demonstrating ex vivo slice generation of conditioned media to acutely incubate with human microglial (“iMGL”) and glutamatergic/ GABAergic neuron tri-cultures and subsequently quantify microglial synaptic engulfment. Ex vivo slices were either saline-injected (“Nontumor”) or glioma xenografted (SU-DIPG-VI, “DMG”), with active optogenetic stimulation (Thy1::ChR2+, active conditioned media, “ACM”) or nonstimulated blue light control (Thy1::ChR2-, conditioned media, “CM”). (B) Representative 3D rendered images from human microglial/glutamatergic/GABAergic neuron tri-cultures exposed to ex vivo slice generated conditioned media. Microglia (white), neurons (blue), PSD95+ excitatory synapses (red), and gephyrin+ inhibitory synapses (green) illustrate microglial synaptic engulfment in tri-culture. Tri-cultures were either treated with conditioned media generated from nonstimulated glioma-bearing slices (“DMG CM”) or from stimulated glioma-bearing slices (“DMG ACM”). Scale bar, 15 um. (C) Volumetric engulfment analysis of microglial engulfment of PSD95+ synapses in tri-culture (“Excitatory”), represented as the % volume of PSD95+ synapse contained in Iba1+ microglia in tri-cultures exposed to ex vivo conditioned media from nonstimulated nontumor slices (“Nontumor CM”) or from stimulated nontumor slices (“Nontumor ACM”). (D) Volumetric engulfment analysis of microglial engulfment of gephyrin+ synapses in tri-culture (“Inhibitory”), represented as the % volume of gephyrin+ synapse contained in Iba1+ microglia in tri-cultures exposed to ex vivo conditioned media from nonstimulated nontumor slices (“Nontumor CM”) or from stimulated nontumor slices (“Nontumor ACM”). (E) Volumetric engulfment analysis of microglial engulfment of PSD95+ synapses in tri-culture (“Excitatory”), represented as the % volume of PSD95+ synapse contained in Iba1+ microglia in tri-cultures exposed to ex vivo conditioned media from nonstimulated glioma-bearing slices (“DMG CM”) or from stimulated glioma-bearing slices (“DMG ACM”). (F) Volumetric engulfment analysis of microglial engulfment of gephyrin+ synapses in tri-culture (“Inhibitory”), represented as the % volume of gephyrin+ synapse contained in Iba1+ microglia in tri-cultures exposed to ex vivo conditioned media from nonstimulated glioma-bearing slices (“DMG CM”) or from stimulated glioma-bearing slices (“DMG ACM”). (G) Ex vivo slice generated conditioned media underwent LC/MS analysis, calculating the ratio of protein abundance (“Protein Abundance Ratio”, y-axis) normalized on total peptide amount, and comparing conditioned media from stimulated glioma-bearing slices (“DMG ACM”) to conditioned media from nonstimulated glioma-bearing slices (“DMG CM”). Individual proteins with abundance ratios >1 are labeled and described. Each individual point represents the average volumetric quantification from images analyzed in one tri-culture (C-F). Data are mean ± s.e.m. One-way ANOVA with Tukey’s post hoc analysis (E); *P<0.05, NS, not significant.

To better understand how glioma-infiltrated brain slice-derived and activity-regulated secreted factors may affect microglial function, we performed an *in vitro* engulfment analysis which quantified microglial synaptic engulfment of excitatory (PSD95) or inhibitory (gephyrin) synapses in the tri-culture system, which enabled investigation of microglia-mediated engulfment of neuron-neuron synapses. We first observed that the mere presence of glioma-infiltrated brain-derived secreted factors – which could be secreted by DMG cells or other cell types in the brain microenvironment influenced by glioma - shifted microglial morphology (**Fig. 4B-C, Supp. Fig. 3A**). When analyzing microglial morphology following exposure to DMG-xenografted brain slice ACM (“DMG ACM”), we observed the formation of elongated, branched microglial ramifications intertwined with neuronal soma and neurites in tri-culture, suggesting extensive interactions of microglia with neurons in response to activity-regulated glioma-associated factors (**Fig. 4B, Supp. Video 2**). These human microglia additionally secreted a variety of proteins in response to treatment with nontumor- or glioma-infiltrated brain-derived factors (**Supp. Fig. 3B-I**).

We observed an activity-regulated increase in excitatory neuron-neuron synapse engulfment by microglia *in vitro* in response to secreted factors from nontumor brain slice ACM (**Fig. 4C**), mirroring the *in vivo* increase in stimulated nontumor models shown above; no changes in inhibitory synaptic engulfment were observed, also in line with the *in vivo* findings described above (**Fig. 4D**). This suggests the presence of an activity-regulated secreted factor or factors that increase excitatory synaptic engulfment by microglia. Upon exploring mass spectrometric analysis of nontumor brain slice secreted proteins between stimulated and nonstimulated CM, we identified activity-regulated candidates VDBP, PICALM, CD13, and APOE (**Supp. Fig 4A-H**) that will be followed up in future work.

Strikingly, there was a substantial downregulation in microglial synaptic engulfment of excitatory neuron-neuron synapses in response to ACM from DMG xenografted brain slices (**Fig. 4E**), suggesting that a secreted factor from the glioma-infiltrated brain suppresses the normal microglial synaptic engulfment of excitatory synapses in response to glutamatergic neuronal activity. Glioma-related influences on inhibitory synaptic engulfment by microglia trended towards a decrease in this assay, but this experiment is presently underpowered (**Fig. 4F**). Upon exploring mass spectrometric analysis of secreted factors from DMG-xenografted brain slices, we found candidate factors upregulated in the CM from optogenetically stimulated brain slices that could potentially drive the observed shift in microglial excitatory synaptic engulfment that include: GST1, PRDX6, and VLIG1 (**Fig. 4G, Supp. Fig. 4A-H**). As expected^13^, we also observed activity-regulated upregulation of shed NLGN3 in glioma-bearing brain slices in comparison with nontumor controls, consistent with our previous findings^13^.

Taken together, these findings suggest that secreted factors from neuron-glioma networks alter microglial synaptic engulfment, leading to an imbalance in excitatory and inhibitory synaptic engulfment in glioma-infiltrated neural circuits.

## DISCUSSION

Neuronal activity promotes glioma progression through a variety of mechanisms, including activity-regulated secretion of paracrine growth factors^10, 12, 13, 96^, direct synaptic signaling between neurons and glioma cells^14–18^, and potassium-evoked glioma currents^15, 17^, while gliomas increase neuronal excitability^15, 19–22^ and functionally remodel^23, 24^ neural networks. Glioma-derived factors like glutamate^19^, as well as synaptogenic proteins such as glypican-3^21^ and thrombospondin-1^23^, promote neuronal hyperexcitability and abundance of excitatory neuron-neuron synapses^21^, alongside a loss in inhibitory synapses^78^, driving heightened excitatory neuronal activity in the glioma microenvironment with consequent tumor cell proliferation and migration in the glioma tumor microenvironment^10, 13, 14, 16–18^. This increased excitatory connectivity and overall circuit activity thereby strengthen the growth and integration of glioma into neuronal networks^23, 24^. However, the contributions of non-neuronal cell types in these malignant neuron-glioma network modifications remain largely unknown.

Here we found that microglia-known mediators of neural circuit refinement^34–44, 46–49^-contribute to this malignant glioma-associated neural circuitry via aberrant synaptic engulfment of excitatory and inhibitory neuron-neuron synapses in the glioma tumor microenvironment. During CNS development and throughout adulthood, microglia precisely refine neural circuits. Microglia are known to utilize GABA-receptive machinery to engulf postnatal inhibitory circuits^38^, while concurrently utilizing complement-receptive pathways to engulf postnatal excitatory synapses^34, 37^. Imbalanced microglial engulfment of excitatory and inhibitory synapses leads to neuronal hyperexcitability^31, 72–76^, increasing seizure susceptibility^31, 38^ and behavioral deficits^31, 38, 79, 80, 82–88^ in rodent models-pathological features which have also been reported in human glioma patients^15, 19–21, 23, 24^. Here we explored microglial synaptic engulfment in glioma-infiltrated brain to better understand how microglia-synaptic interactions may change in a cancer-specific manner.

We identified activity-regulated mechanisms of microglia-mediated excitatory synaptic engulfment in both homeostatic and oncogenic contexts. Microglia in the healthy, nontumor-infiltrated brain respond to optogenetically stimulated glutamatergic neuronal activity by engulfing more excitatory synapses, whereas microglia in the malignant glioma-infiltrated brain apparently lose this normal mechanism of feedback regulation and do not increase excitatory engulfment in response to glutamatergic neuronal activity. Shedding light on this dysregulation, we found that a normal activity-regulated increase in complement pathway gene expression is lost in glioma-associated microglia. Remarkably, instead of the feedback control of excitability through engulfment of excitatory synapses found in the healthy cortex, glioma-associated microglia aberrantly and excessively engulf inhibitory synapses in response to glutamatergic neuronal activity. This inversion of healthy physiology is mediated by glioma-secreted or glioma-regulated secreted factors from other tumor-associated neural cells, as conditioned medium from glioma-xenografted brain slices recapitulates this microglial behavior. Taken together, these findings indicate that microglia become reprogrammed in glioma-infiltrated neural networks to specifically downregulate machinery which would otherwise establish feedback regulatory control of circuit excitability through microglial engulfment of excitatory synapses.

## Limitations of the Study

The secreted factors mediating the activity-regulated increase in excitatory synaptic engulfment in the healthy brain remain to be identified, as do the secreted factors that invert this effect and instead promote elevated microglial inhibitory synapse engulfment in the context of glioma. Furthermore, it remains an open question how glioma-associated microglial synaptic engulfment differentially affects excitatory synapses on excitatory neurons versus those on inhibitory neurons, and similarly how glioma-associated microglia could differentially affect inhibitory synaptic engulfment on excitatory versus inhibitory neurons. Furthermore, both glutamatergic^15^ and (depolarizing) GABAergic^16^ neuron-to-glioma synapses form on DMG cells; it is not yet known how microglia interact with these malignant synapses. Future studies should identify the microglial subpopulation(s) that aberrantly engulf inhibitory synapses in DMG and determine if microglia differentially engulf neuron-to-neuron versus neuron-to-glioma synapses. Finally, while we have shown that DMG cells induce hyperexcitability and also alter the proportion of excitatory vs inhibitory synapses engulfed in response to glutamatergic activity, we have not shown directly the extent to which the altered excitatory:inhibitory balance of microglial synaptic engulfment contributes to this hyperexcitable circuit pathophysiology in DMG. When the molecular mechanisms mediating this are identified in future work, such direct demonstration will be possible.

## Conclusions

Just as glioma reprograms microglia into an immunosuppressive state that counters anti-tumor immunity^56, 68, 70, 71^, we demonstrate here that glioma also reprograms microglial-synaptic interactions to promote malignant neural circuit physiology. Future studies will further elucidate the paracrine signaling factors that mediate this microglial dysregulation.

## ACKNOWLEDGEMENTS

We express our gratitude to the children and their families who generously donated tumor tissue for research. This work was supported by grants from the National Institute of Neurological Disorders and Stroke (R01NS092597 to M.M.), NIH Director’s Pioneer Award (DP1NS111132 to M.M.), National Cancer Institute (P50CA165962, R01CA258384, U19CA264504 to M.M.), Gatsby Charitable Foundation (Gatsby Initiative in Brain Development and Psychiatry, to M.M.), Oscar’s Kids Foundation (to M.M.), McKenna Claire Foundation (to M.M.), Kyle O’Connell Foundation (to M.M.), Yuvaan Tiwari Foundation (to M.M.), Virginia and D.K. Ludwig Fund for Cancer Research (to M.M.), Waxman Family Research Fund (to M.M.), Will Irwin Research Fund of the Pediatric Cancer Research Foundation (to M.M.), Austin Strong Foundation (M.M.), Avery Huffman DIPG Foundation (to M.M.), Chadtough Defeat DIPG (to M.M.), Jacob Van de Roovaart Glioblastoma Research Fund (to M.M.), the Chambers-Okamura Endowed Directorship for Pediatric Neuro-Immuno-Oncology (MM), Botha-Chan Family Research Fund (to M.M.).

The authors thank Dr. Won-Suk Chung and Dr. Joon-Hyuk Lee from Korea Advanced Institute of Science & Technology (KAIST) for generously providing fluorescent synaptic AAVs used in our study, and Dr. Michael Dolan from Trinity College Dublin and Alex’s Lemonade Stand’s Childhood Cancer Data Lab-including Dr. Jaclyn Taroni, Dr. Joshua Shapiro, and Dr. Ally Hawkins-for their expert sequencing advising.

## RESOURCE AVAILABILITY

### Lead contact

Further information and requests for resources and reagents should be directed to and will be fulfilled by the lead contact, Michelle Monje.

### Materials availability

This study did not generate new unique reagents.

### Data and code availability

All single-nucleus RNA sequencing datasets will be deposited to GEO.

## METHODS

### KEY RESOURCES TABLE

| REAGENT or RESOURCE | SOURCE | IDENTIFIER |
| --- | --- | --- |
| **Antibodies** |  |  |
| Anti-rabbit IBA1 | Wako Chemicals | Cat#019-19741;<br>RRID: AB_839504 |
| Anti-mouse human nuclear antigen 235-1 | Millipore | MAB1281 |
| Anti-chicken GFP | Abcam | ab13970 |
| Anti-rabbit Ki67 | Abcam | ab15580 |
| Alexa Fluor 488 donkey anti-rabbit | Jackson ImmunoResearch | 711-545-152 |
| Alexa Fluor 488 donkey anti-chicken | Jackson ImmunoResearch | 703-545-155 |
| Alexa Fluor 555 donkey anti-rabbit | Jackson ImmunoResearch | 711-565-152 |
| Alexa Fluor 594 donkey anti-chicken | Jackson ImmunoResearch | 711-585-152 |
| Alexa Fluor 647 donkey anti-rabbit | Jackson ImmunoResearch | 711-605-152<br>RRID: AB_2492288 |
| DAPI | Thermo Fisher Scientific | Cat# 62247; |
| Prolong Gold Mounting Medium | Life Technologies | Cat# P36930; |
| Normal Donkey Serum | Jackson Immunotherapy | Cat# 017-000-121 |
| **Bacterial and virus strains** |  |  |
| hSyn-PSD95-mCherry-eGFP (ExPost) | Dr. Won-Suk Chung, KAIST | N/A |
| hSyn-gephyrin-mCherry-eGFP (InhiPost) | Dr. Won-Suk Chung, KAIST | N/A |
| **Biological samples** |  |  |
| **Chemicals, peptides, and recombinant proteins** |  |  |
| 8% Glutaraldehyde | Electron Microscopy Services | Cat#16000 |
| 16% Paraformaldehyde | Electron Microscopy Services | Cat#15700 |
| 0.1 M sodium cacodylate | Electron Microscopy Services | Cat#12300 |
| DMEM | Thermo Fisher Scientific | 11320082 |
| Neurobasal-A | Invitrogen | 10888-022 |
| B27-A | Thermo Fisher Scientific | 12587-010 |
| Heparin solution | STEMCELL Technologies | 07980 |
| Human bFGF | Shenandoah Biotech | 100-146 |
| Human bEGF | Shenandoah Biotech | 100-26 |
| Human PDGF-AA | Shenandoah Biotech | 100-16 |
| Human PDGF-BB | Shenandoah Biotech | 100-18 |
| GDNF | Shenandoah Biotech | 200-37 |
| BDNF | Shenandoah Biotech | 100-01 |
| Insulin | Sigma-Aldrich | I9278 |
| 2-Mercaptoethanol | Sigma-Aldrich | 63689 |
| TrypLE Express Enzyme | Invitrogen | 12604013 |
| Laminin Mouse Protein | Thermo Fisher Scientific | 23017015 |
| Poly-L-Lysine solution | Sigma-Aldrich | P4707 |
| BrainPhys Neuronal medium | STEMCELL Technologies | 05790 |
| Glutamax | Invitrogen | 35050-061 |
| Sodium pyruvate | Invitrogen | 11360070 |
| MEM non-essential amino acids | Thermo Fisher Scientific | 11140076 |
| Antibiotic-antimycotic | Thermo Fisher Scientific | 15240096 |
| 5-ethynyl-2'-deoxyuridine | Invitrogen | E10187 |
| Optimum cutting temperature (OCT) | Tissue-Tek | 4583 |
| Triton X-100 | Sigma-Aldrich | 9036-19-5 |
| μ-Plate 24 Well Black ID 14 mm | Ibidi | 82421 |
| MEM Non-Essential Amino Acids Solution (100X) | Thermo Scientific | 11140050 |
| Advanced DMEM/F-12 | Thermo Scientific | 12634010 |
| Doxycycline hyclate, ≥93.5% (HPLC), Powder | Sigma-Aldrich | D9891-1G |
| Recombinant Human BDNF Protein | R&D Systems | 248-BDB-005 |
| Recombinant Human NT-3 Protein | R&D Systems | 267-N3-005 |
| STD Human Rec Insulin 5 mg/ml 20ml | Lonza | BE02-033E20 |
| Recombinant Human IL-34 (carrier-free) 10 μg | BioLegend | 577902 |
| Recombinant Human IL-6, 20.9 kDa | PeproTech | 200-06 |
| Recombinant Human IL-3 | PeproTech | 200-03 |
| Recombinant Human M-CSF | PeproTech | 300-25 |
| Recombinant Human TGF-β1 (HEK293 derived) | PeproTech | 100-21 |
| Recombinant VEGF Human Protein | Thermo Scientific | PHC9394 |
| Insulin-Transferrin-Selenium (ITS -G) (100X) | Thermo Scientific | 41400045 |
| HEPES | Gibco | 15630080 |
| GlutaMAX | Gibco | 35050061 |
| Penicillin- Streptomycin | Gibco | 15070063 |
| Poly-D-lysine | Corning | 354210 |
| DMEM | Gibco | 11960-044 |
| Tris pH 8.0 | Invitrogen | AM9856 |
| Sucrose | Sigma-Aldrich | S0389-5KG |
| KCl | Invitrogen | AM9640G |
| MgCl<sub>2</sub> | Invitrogen | AM9530G |
| RNasin Plus | Promega | N2611 |
| Protease Inhibitor Cocktail, 50X | Promega | G652A |
| DTT | New England Biolabs | 7016L |
| Actinomycin D | Millipore Sigma | A9415 |
| Triptolide | Millipore Sigma | 645900 |
| Anisomycin | Millipore Sigma | A9789 |
| Nuclease free BSA | Millipore Sigma | 126609 |
| **Critical commercial assays** |  |  |
| PhiX | Illumina | FC-110-3002 |
| 10X Chromium Next GEM Single Cell 3' HT Kits v.3.1 | 10X Genomics | 1000348 |
| Click-iT EdU cell proliferation kit | Invitrogen | C10339 |
| **Deposited data** |  |  |
| Mouse brain single nucseq | Gene Expression Omnibus | TBD |
| **Experimental models: Cell lines** |  |  |
| H3K27M-MADR | This paper | N/A |
| SU-DIPG-VI | Stanford Medical Center | N/A |
| ioMicroglia | Bit.bio | io1029S |
| ioGlutamatergic Neurons | Bit.bio | io1001S |
| ioGABAergic Neurons | Bit.bio | io1003S |
| **Experimental models: Organisms/strains** |  |  |
| NOD.Cg-*Prkdc*<sup>scid</sup> *Il2rg*<sup>tm1Wjl</sup>/SzJ | The Jackson Laboratory | Strain #:005557 |
| B6.SJL-*Ptprca* *Pepcb*/BoyJ | The Jackson Laboratory | Strain #: 002014 |
| C57BL/6J | The Jackson Laboratory | Strain #: 000664 |
| **Oligonucleotides** |  |  |
| **Recombinant DNA** |  |  |
| **Software and algorithms** |  |  |
| FIJI | NIH | https://imagej.net/software/fiji/ |
| Adobe Illustrator 2024 V28.5 | Adobe | N/A |
| Imaris V10.0.0 | Oxford Instruments | N/A |
| Zen | Zeiss | https://www.zeiss.com/microscopy/us/products/microscope-software/zen.html |
| Graphpad Prism | Dotmatics | https://www.graphpad.com/ |
| **Other** |  |  |
| Zeiss LSM980 Confocal Microscope | Zeiss | 980 |
| Leica VT 1200S Vibratome | Leica | 1200S |
| Leica HM450 Microtome | Leica | HM450 |
| FACSAria II | BD Biosciences | SORP |
| FACS Symphony A5 | BD Biosciences | SORP |
| Fiber Optic Rotary Joint Patch Cables (Optogenetics) | Thorlabs | RJPSL2 |
| Mono Fiberoptic Cannula_(Optogenetics) | Doric Lenses Inc. | MFC_200/240-0.22_4.5mm_MF2.5_FLT |
| Optofluid Cannula with interchangeable injectors (Optogenetics) | Doric Lenses Inc. | iOFC_M3_250/350_4.0mm_PLG G |
| 10X Chromium X | 10X Genomics | 1000331 |
| Sony MA900 Sorter | Sony Biotechnology | LE-MA900FP |
| Illumina NextSeq2000 | Illumina | 20038897 |

### STAR Methods

#### EXPERIMENTAL METHOD AND STUDY PARTICIPANT DETAILS

##### Animal Models

All animal experiments were conducted in accordance with protocols approved by the Stanford University Institutional Animal Care and Use Committee (IACUC) and performed in accordance with institutional guidelines. Animals were housed according to standard guidelines with unlimited access to water and food, under a 12 h light: 12 h dark cycle, a temperature of 21 °C and 60% humidity. For brain tumour allograft or xenograft experiments, the IACUC has a limit on indications of morbidity (as opposed to tumour volume). Under no circumstances did any of the experiments exceed the limits indicated and mice were immediately euthanized if they exhibited signs of neurological morbidity or if they lost 15% or more of their initial body weight.

##### Patient-Derived Diffuse Midline Glioma Cells

Diffuse midline glioma cultures were established from patient-derived specimens with informed consent, following a protocol approved by the Stanford University Institutional Review Board (IRB). For all tumor studies, Throughout the culture period, all cultures were subjected to monitoring for authenticity via short tandem repeat (STR) fingerprinting and routine mycoplasma testing was conducted. Characteristics of the glioma models used (patient information, molecular characterization, and other characteristics) have been previously reported ^10, 91, 97^. The glioma cultures were cultivated as neurospheres in serum-free medium composed of DMEM (Invitrogen), Neurobasal(-A) (Invitrogen), B27(-A) (Invitrogen), heparin (2 ng ml−1), human-bFGF (20 ng ml−1) (Shenandoah Biotech), human-bEGF (20 ng ml−1) (Shenandoah Biotech), human-PDGF-AA (10 ng ml−1) (Shenandoah Biotech), and human-PDGF-BB (10 ng ml−1) (Shenandoah Biotech). The neurospheres were dissociated using TrypLE (Gibco) for seeding or to generate single cell suspensions for xenografting (see ‘***Orthotopic xenografting’***).

##### Stereotaxic Surgery and Viral Vectors

Mice were anesthetized with 1-4% isoflurane and placed in a stereotaxic apparatus. Expression of fluorescently-labeled PSD95 or gephyrin was achieved by injecting 1 µl of AAV9-CW3SL::hSyn-PSD95-mCherry-eGFP ^92^or AAV9-CW3SL::hSyn-gephyrin-mCherry-eGFP ^92^(virus titer= 1.0×10^11^ gc/ml), generously provided by the laboratory of Won-Suk Chung at KAIST. Both viral vectors were packaged into AAV9 capsids at the Stanford University Gene Vector and Virus Core. Viral vectors were unilaterally injected using Hamilton Neurosyringe and Stoelting stereotaxic injector over 5 minutes. Cortical (M2) coordinates were: 1 mm anterior to bregma, 0.5 mm lateral to midline, and 1 mm deep to cranial surface.

##### Generation of H3.3 K27M MADR cells

The H3.3 K27M MADR tumor cell cultures were generated using the same technique as previously described ^95^. Briefly, Gt(ROSA)26Sortm4(ACTB-tdTomato,-EGFP)Luo/J and Gt(ROSA)26Sortm1.1(CAG-EGFP)Fsh/Mmjax mice (JAX Mice) were crossed with wild-type CD1 mice (Charles River) to produce heterozygous mice. Male and female embryos between E12.5 and E15.5 were subjected to *in utero* electroporations. Pregnant dams were individually housed, and pups remained with their mothers until P21 in the institutional animal facility (Stanford University). The MADR tumor cell line used here was generated by dissociating and sorting GFP+ tumor cells from female heterozygous mTmG mice. Subsequently, MADR cultures were maintained as neurospheres in serum-free medium composed of DMEM (Invitrogen), Neurobasal(-A) (Invitrogen), B27(-A) (Invitrogen), heparin (2 ng ml−1), human bFGF (20 ng ml−1) (Shenandoah Biotech), human bEGF (20 ng ml−1) (Shenandoah Biotech), human PDGF-AA (10 ng ml−1) (Shenandoah Biotech), insulin (Sigma-Aldrich), and 2-mercaptoethanol (Sigma-Aldrich). The neurospheres were dissociated using TrypLE (Gibco) for seeding or to generate single cell suspensions for allografting (see ‘***Orthotopic allografting***).

##### Orthotopic xenografting

For all xenograft studies, NSG mice (NOD-SCID-IL2R gamma chain-deficient, The Jackson Laboratory) were used. Male and female mice were used equally. For all synaptic engulfment studies, a single-cell suspension from patient-derived DIPG culture SU-DIPG-VI was stereotactically injected into premotor (M2) frontal cortex. A single-cell suspension of all cultures was prepared in sterile PBS (see ‘***Patient-Derived Diffuse Midline Glioma Cells***’) immediately before surgery. Animals at postnatal day (P) 28–P35 were anaesthetized with 1–4% isoflurane and placed on stereotactic apparatus. Under sterile conditions, the cranium was exposed via a midline incision and a 31-gauge burr hole made at exact coordinates. For cortical injections coordinates were: 1 mm anterior to bregma, 0.5 mm lateral to midline, and 1 mm deep to cranial surface. Approximately 300,000 cells in 3uL sterile PBS were injected using a 31-gauge Hamilton syringe at an infusion rate of 0.4 µl min−1 with a digital pump. At completion of infusion, the syringe needle was allowed to remain in place for a minimum of 2 min to minimize backflow of the injected cell suspension, then manually withdrawn. The wound was closed using sterile sutures and treated with Neo-Predef with Tetracaine Powder. DIPG xenografted mice were euthanized 8 weeks following DIPG cell injection.

##### Orthotopic allografting

CD1 mice (the Jackson Laboratory) were crossed with C57BL/6 J mice (the Jackson Laboratory) to mimic the genetic background of the mouse which generated the H3.3K27M MADR tumor cell line (see ‘***Generation of H3.3 K27M MADR cells’***). For all allograft studies, CD1xBL6 mice were used. Male and female mice were used equally. For all sequencing studies, a single-cell suspension from murine MADR culture (H3.3 K27 MADR line 1) was stereotactically injected into premotor (M2) frontal cortex. A single-cell suspension of all cultures was prepared in sterile PBS (see ‘***Generation of H3.3 K27M MADR cells’***) immediately before surgery. Animals at postnatal day (P) 28–P35 were anaesthetized with 1–4% isoflurane and placed on stereotactic apparatus. Under sterile conditions, the cranium was exposed via a midline incision and a 31-gauge burr hole made at exact coordinates. For cortical injections coordinates were: 1 mm anterior to bregma, 0.5 mm lateral to midline, and 1 mm deep to cranial surface. Approximately 200,000 cells in 3uL sterile PBS were injected using a 31-gauge Hamilton syringe at an infusion rate of 0.4 µl min−1 with a digital pump. At completion of infusion, the syringe needle was allowed to remain in place for a minimum of 2 min to minimize backflow of the injected cell suspension, then manually withdrawn. The wound was closed using sterile sutures and treated with Neo-Predef with Tetracaine Powder. MADR allografted mice were euthanized 5 weeks following MADR cell injection.

##### Fibre-optic placement and in vivo optogenetic stimulation

For optogenetics experiments in DIPG xenografted mice, Thy1::ChR2+/− mice (line 18, The Jackson Laboratory, C57BL/6 J background) were crossed with NSG mice (NOD-SCID-IL2R gamma chain-deficient, The Jackson Laboratory) to produce the Thy1::ChR2;NSG genotype.

For optogenetics experiments in MADR allografted mice, Thy1::ChR2+/− mice (line 18, The Jackson Laboratory, C57BL/6 J background) were crossed with CD1 mice (the Jackson Laboratory) to produce the Thy1::ChR2;CD1/BL6 genotype.

Experiments for the acute neuronal optogenetic stimulation paradigm of glioma xenografts were performed as previously described ^10, 13, 14^. In brief, a fibre-optic ferrule (Doric Lenses) was placed at M2 of the right hemisphere with the following coordinates: 1.0 mm anterior to bregma, 0.5 mm lateral to midline, −0.7 mm deep to the cranial surface at four weeks post DIPG xenograft or three weeks post MADR allograft. Following 4 weeks recovery for DIPG xenografted cohorts and 1 week recovery for MADR allografted cohorts, all mice were connected to a 473 nm diode-pumped solid-state (DPSS) laser system via a mono fibre patch cord. In awake mice, pulses of light with a power measured at 10 mW from the tip of the optic fiber (200 µm core diameter, NA=0.22 - Doric lenses) were administered at a frequency of 20 Hz for a period of 30 s, followed by 90 s recovery in a repeated cycle for a single session of 30 minutes. This power represents approximately 30 mW cm−2 light density at the tip of the patch cord; with the optical ferrule placed just below the pial surface this would deliver 3 mW cm−2 approximately midway through the cortex ^93, 98^. The mice were monitored for their unidirectional ambulation response to light stimulation, confirming correct ferrule placement over M2 of the right hemisphere and effective neuronal stimulation. Animals confirmed as ChR2-negative had no response to light stimulation. Mice were euthanized 24 hours following the stimulation session.

##### Cerebral Slice Conditioned Media

Thy1::ChR2- or Thy1::ChR2+ mice aged 4 weeks (P28 to P30) were utilized to collect conditioned media from activated excitatory cortical neurons. Mice had either been previously saline-injected or xenografted with DIPG in the cortical M2 region (see ‘***Orthotopic xenografting***’). Brief exposure to isoflurane induced unconsciousness in the mice before immediate decapitation. Extracted brains (cerebrum) were inverted and placed in an oxygenated sucrose cutting solution, then sliced at 300μm to target the cortical region (M2 area). Slices (n=4 per mouse) were transferred to ACSF and allowed to recover for 30 minutes at 37°C, followed by an additional 30 minutes at room temperature. After recovery, the slices were transferred to fresh ACSF and positioned under a blue-light LED using a microscope objective. The optogenetic stimulation paradigm consisted of 20-Hz pulses of red light for 30 seconds on, followed by 90 seconds off, repeated over a period of 30 minutes. Conditioned medium from the surrounding area was collected and stored frozen at −80°C. Stimulated slices were postfixed in 4% paraformaldehyde (PFA) for 30 minutes before cryoprotection in 30% sucrose solution for 48 hours.

##### Human iPSC-derived Cells

Human iPSC-derived microglia (io1029S, bit.bio), GABAergic neurons (io1003S, bit.bio) and glutamatergic neurons (io1001S, bit.bio) were thawed according to manufacturer’s specifications. In short, GABAergic and glutamatergic neuron co-culture was initiated on day 0, utilizing an growth media compatible for both neuronal subtypes, and maintaining co-cultures for 14 days according to manufacturer’s recommendations. Concurrently, microglial cultures were initiated on day 0, utilizing a growth media and maintenance protocol according to manufacturer’s recommendations.

On day 14, microglia were dissociated and plated with the GABAergic/glutamatergic co-culture using growth media compatible to all 3 cell types in line with manufacturer’s recommendations. Microglia, GABAergic neurons, and glutamatergic neurons were incubated in culture for 24 hours before beginning further experimental procedures. Human ioMicroglia, ioGABAergic, and ioGlutamatergic neurons have been extensively characterized and functionally validated in manufacturer workflows.

##### Magnetic resonance imaging

Anatomic magnetic resonance imaging (MRI) images of MADR glioma allografts were acquired with a T2-weighted Turbo RARE pulse sequence with fat suppression and the parameters TE = 33ms, TR = 2500 ms, RARE factor = 8, 4 averages, 19 slices, slice thickness 0.7 mm, field of view = 25 mm x 25 mm and image matrix 512 x 256.

#### METHOD DETAILS

##### Immunohistochemistry

For immunohistochemical analyses, mice were transcardially perfused with 20 mL of 0.1M phosphate buffer saline (PBS) followed by 20 mL of 4% paraformaldehyde (PFA), postfixed overnight in 4% PFA at 4°C, then transferred to 30% sucrose for 24 hours. Brains were flash frozen, embedded in optimum cutting temperature (OCT; Tissue-Tek) and sectioned coronally in 40um increments on a temperature-controlled sliding microtome (Leica, HM450).

Coronal sections in the specific region of interest (for example, M2) were collected and stained with antibodies following an incubation in blocking solution (3% normal donkey serum, 0.3% Triton X-100 in tris buffer saline, TBS) at room temperature for 2 hours. For synaptic engulfment analysis, rabbit anti-IBA1 (FUJIFILM Wako 019-19741, 1:500) and mouse anti-HNA (abcam ab191181, 1:250) were diluted in 1% blocking solution (1% normal donkey serum in 0.3% Triton X-100 in TBS) and incubated overnight at 4°C. For tumor histology, brain sections were first stained using the Click-iT EdU cell proliferation kit (Invitrogen, C10339 or C10337) according to the manufacturer’s protocol, then chicken anti-GFP (abcam ab13970, 1:500) and rabbit anti-Ki67 (abcam ab15580, 1:250) were diluted in 1% blocking solution and incubated overnight at 4°C. 24 hours following primary antibody incubation, sections were rinsed 3 times in 1x TBS and incubated in secondary antibody solution for 2 hours at room temperature. All secondary antibodies were used at 1:500 concentration including Alexa 488 anti-rabbit, Alexa 488 anti-mouse, Alexa 488 anti-chicken, Alexa 594 anti-chicken, Alexa 647 anti-goat, Alexa 647 anti-rat, Alexa 647 anti-rabbit, Alexa 405 anti-guinea pig, and Alexa 555 anti-rabbit (all Jackson ImmunoResearch). Sections were then rinsed three times in 1x TBS and mounted with ProLong Gold (Life Technologies, P36930).

##### Confocal Microscopy

All images analyses were performed by experimenters blinded to the experimental conditions or genotype. Images were captured by acquiring Airyscan z-stacks using a Zeiss LSM980 scanning confocal microscopy. For each animal, 3-5 regions of interest were imaged using 0.2 µm z-steps.

##### 3D-rendering and Quantification

Following Airyscan image processing, Imaris was used to create 3D volume surface renderings of each z-stack. Surface rendered images were used to determine the volume of the microglia, glioma cells, and all synaptic structures. To determine % engulfment, the following calculation was used: Volume of internalized synapse (µm^3^)/Volume microglial cell (µm^3^), following previously-established protocols in microglial synaptic engulfment analyses ^34, 37^.

##### Protein Identification by Mass Spectrometry

###### Sample preparation

Media samples were concentrated to 4 mg/mL using spin columns with a 3 KDa molecular weight cut-off (MWCO), then buffer-exchanged into 50 mM ammonium bicarbonate. Protein concentration was determined using the Bio-Rad Protein Assay Kit II (#500-0002) according to the manufacturer’s instructions. For each sample, 30 µg of protein was used. DTT was added to a final concentration of 10 mM, and samples were incubated at 60 °C for 30 minutes, followed by cooling to room temperature. Iodoacetamide was then added to a final concentration of 10 mM, and samples were incubated in the dark at room temperature for 30 minutes. Proteins were subsequently digested overnight at 37 °C using Trypsin/Lys-C Mix, Mass Spec Grade (Promega).

###### Albumin Removal

Albumin was selectively depleted from the sample using an antibody-based affinity column engineered for high-specificity binding to serum albumin. Media samples were first concentrated, then centrifuged at 14,000 rpm for 15 minutes at 4 °C. The resulting supernatant was collected and passed through 1 mL Immobilized Anti-BSA Antibody columns (G-BioSciences) three times, followed by a wash step. The flow-through, containing non-BSA proteins, was collected and concentrated to 4 mg/mL using 3 kDa MWCO spin columns, then buffer-exchanged into 50 mM ammonium bicarbonate.

###### nanoLC-MS/MS

NanoLC separation was performed using a Thermo Fisher Ultimate 3000 system (Milford, MA). Mobile phases consisted of solvent A: 0.1% formic acid (v/v) in water, and solvent B: 0.1% formic acid (v/v) in 90% acetonitrile. Tryptic peptides were first loaded onto a µPrecolumn Cartridge, followed by separation on a reverse-phase analytical HPLC column (Thermo Fisher Scientific, ES900). The chromatographic gradient was as follows: 2% to 10% solvent B over 10 minutes, 10% to 30% B over 85 minutes, followed by a wash at 99% B for 10 minutes and re-equilibration at 4% B for 5 minutes. Mass spectrometry was conducted on an Orbitrap Exploris (Thermo Fisher Scientific). MS1 scans were acquired at 80,000 resolution over a scan range of m/z 300–2000, with 100% AGC target and maximum injection time set to auto. MS/MS scans were performed at 30,000 resolution with a 200% AGC target, auto maximum injection time, 30% normalized HCD collision energy, and an intensity threshold of 1.0e4.

###### Database search

Data analysis was conducted using Proteome Discoverer 2.5 (Thermo Fisher Scientific). MS and MS/MS spectra were searched against the SwissProt Human or Mouse database, containing all protein entries as of March 2025. Searches were performed using the Sequest HT algorithm with the following parameters: enzyme specificity set to Trypsin, minimum peptide length of 5, maximum peptide length of 144, precursor mass tolerance of 10 ppm, and fragment mass tolerance of 0.02 Da. Dynamic modifications included oxidation of methionine (Oxidation-M), N-terminal acetylation, and N-terminal methionine loss. Carbamidomethylation of cysteine (Carbamidomethyl-C) was set as a static modification. Label-free quantification of individual and grouped protein abundances was performed using the Minora Feature Detector. For both the processing and consensus workflows, results were filtered to a maximum false discovery rate (FDR) of 1%. Protein abundance values and ratios were normalized based on total peptide amount. Protein abundance values and ratios were normalized on total peptide amount. nanoLC-MS/MS and data analysis was performed by Applied Biomics (www.appliedbiomics.com/)

##### EdU Incorporation Assay

EdU staining of glioma monocultures was performed on glass coverslips in 96-well plates which were pre-coated with poly-l-lysine (Sigma) and laminin (Thermo Fisher Scientific). Neurosphere cultures were dissociated with TrypLE and plated onto coated coverslips with growth factor-depleted medium. After 24 h the cells were fixed with 4% PFA in PBS for 20 min and then stained using the Click-iT EdU kit and protocol (Invitrogen) and mounted using Prolong Gold mounting medium (Life Technologies). Confocal images were acquired on a Zeiss LSM980 using Zen 2011 v8.1. Proliferation index was determined by quantifying the fraction of EdU-labelled cells divided by HNA-labeled cells using confocal microscopy at 20× magnification. Quantification of images was performed by a blinded investigator.

##### Slice preparation for electrophysiology

Coronal brain slices (300 µm) containing the hippocampus were prepared from 6–8-week-old mice of both sexes using a vibratome (Leica VT 1200S, Leica, Germany). All procedures followed a protocol approved by the Stanford University Administrative Panel on Laboratory Animal Care (APLAC). Briefly, mice were anesthetized with isoflurane before brain extraction. Following decapitation, brains were rapidly removed and placed into ice-cold (0-4°C) artificial cerebrospinal fluid (ACSF) composed of (in mM): 110 choline chloride, 2.5 KCl, 1.2 Na2HPO4, 3.5 MgCl2, 0.5 CaCl2, 1.2 MgSO4, 10 glucose, and 26 NaHCO3, bubbled with 95% O2 and 5% CO2 to achieve a pH of 7.3 (osmolality: ∼305 mOsm). Slices were then incubated for 15–20 minutes at 34°C in ACSF containing (in mM): 126 NaCl, 2.5 KCl, 1.2 Na2HPO4, 1 MgCl2, 1.5 CaCl2, 10 glucose, and 26 NaHCO3, with the pH adjusted to 7.3 (osmolality: ∼305 mOsm). This step was followed by a one-hour incubation at room temperature (approximately 25°C) to allow for cellular equilibration before transferring to the recording chamber. All recordings were conducted at the physiologically optimal temperature of 34°C.

##### Electrophysiology

Brain slices were placed in a recording chamber mounted on an upright microscope (BX51WI, Scientifica, USA) and continuously perfused with oxygenated artificial cerebrospinal fluid (ACSF; 95% O₂ / 5% CO₂) at a flow rate of 2–3 mL/min. All experiments were conducted at or near physiological temperature (33–34°C). To investigate the increased excitability of excitatory neurons within the hippocampus, pyramidal cells were recorded in the CA1 region of the stratum pyramidale (Fig.) of the hippocampus. Pyramidal cells were identified and visualized using differential interference contrast (DIC), infrared optics and an sCMOS camera (Kinetix, Teledyne Vision Solutions©, USA).

##### Postsynaptic current recordings

Spontaneous excitatory postsynaptic currents (sEPSCs) and inhibitory postsynaptic currents (sIPSCs) were recorded from pyramidal cells in standard ACSF using whole-cell voltage-clamp mode. Borosilicate glass pipettes (5–8 MΩ) were filled with an internal solution containing (in mM): 9 CsCl, 123 CsMeSO3, 10 HEPES, 2 Mg-ATP, 0.3 Na-GTP, 5 Na-phosphocreatine, and 1 EGTA (pH 7.35, 295 mOsm). sEPSCs were recorded at a holding voltage of –70 mV, a potential at which no net currents occur through GABAA receptors. sIPSCs were isolated by recording at a holding potential of 0 mV, the reversal potential of AMPA- and NMDA-receptor-mediated currents. Recordings were excluded if the holding current exceeded –50 pA or if the series resistance increased by more than 30 MΩ, which was continuously monitored. Since neuronal excitability correlates with an increase in field potential amplitude, field excitatory postsynaptic potentials (fEPSPs) were recorded using a glass pipette (1 MΩ) filled with 1 M NaCl, positioned within the CA1 region. Presynaptic stimulation was performed using a concentric bipolar microelectrode (FHC Microelectrodes, USA), inserted approximately 20–50 µm deep in the CA3 region (Schaffer collateral) of the hippocampus. Before each recording session, the location and intensity of the stimulus were adjusted to ensure consistent inward currents of similar amplitude were evoked. Input currents were delivered through an ISO-Flex stimulus isolator (AMPI, IL) with precise control over the duration of the TTL pulse (a single square wave lasting 1 ms) generated by a Digidata 1550B system, controlled via Clampex software. A series of pulses, ranging in intensity from 0 to 100 µA, were delivered at 10-second intervals to measure the resulting output fEPSPs. To enhance clarity, stimulus artifacts were removed from the field potential recordings. All electrophysiological data were collected using a MultiClamp 700B amplifier, filtered at 2 kHz, digitized at 20 kHz with the Digidata 1550B system, and analyzed using Clampex 11 software (Molecular Devices: Axon Instruments, USA).

##### Single nuclei sequencing

###### Single cell nuclei isolation and sorting

Nuclei were isolated and prepared for sorting in a BSL-2 tissue culture hood. All tissue preparation steps were performed on ice. Mice were perfused using perfusion buffer containing 5 µg ml^−1^ of actinomycin D and 10 µM triptolide diluted in UltraPure DNase/RNase-free distilled water. Tissue was microdissected and punched using Miltex® Biopsy Punch w/ Plunger, 1.0mm (Integra, 33-31AA-P/25) and visually inspected to confirm cortical or striatal anatomy. Punches were subsequently added to their respective Wheaton Dounce homogenizer containing 1 mL of ice-cold nuclei isolation medium (10 mM Tris pH 8.0, 250 mM sucrose, 25 mM KCl, 5 mM MgCl2, 0.1% Triton X-100, 1% RNasin Plus, 1x protease inhibitor, 0.1 mM DTT, 5 µg ml^−1^ of actinomycin D, 10 µM triptolide and anisomycin in UltraPure DNase/RNase-free distilled water). Tissues were mechanically dissociated by ten strokes with the loose Dounce pestle followed by ten strokes with the tight pestle. Homogenates were passed through a 70 μm cell strainer and centrifuged at 900*g* for 15 minutes to pellet the nuclei. Nuclei were resuspended in 1 mL of resuspension buffer (1× phosphate buffer saline (PBS), 1% nuclease free BSA and 0.5% RNasin Plus) and centrifuged at 900*g* for another 15 minutes to pellet the nuclei. Supernatant was removed and nuclei were then stained by adding 1 mL of resuspension buffer containing 0.1 µg mL^−1^ of DAPI. Afterwards, nuclei were filtered through a 40 μm cell strainer into FACS tubes immediately prior to FACS sorting using a Sony MA900 sorter with a 70 µm chip. A standard gating strategy was applied to all samples. First, nuclei were gated on their size and scatter properties. Next, doublet discrimination gates were used to exclude aggregates of nuclei. Lastly, nuclei were gated on DAPI. Single nuclei were sorted into chilled PCR tubes containing 10 µL of resuspension buffer. Nuclei were verified under microscope where counts were verified using a Fuchs-Rosenthal counting chamber.

###### RNA-seq library construction and sequencing

Libraries were constructed using 10X Chromium Next GEM Single Cell 3′ HT Kits v.3.1. After sorting and in accordance with the manufacturer’s instructions, nuclei were carefully handled and promptly added to the master mix after which they were loaded onto a 10X Chromium Next GEM Chip M and ran on a 10X Chromium X instrument with a target recovery of 20,000 nuclei per sample. In alignment with the manufacturer’s protocol, we generated 10X 3′ RNA-seq libraries with subsequent production of complementary DNA amplification and library construction. Quality controls were performed at each appropriate step to ensure sample quality. Generated libraries were loaded at 650 pM along with 1% PhiX control on an Illumina NextSeq2000 and paired-end sequenced (28 cycles read1, 10 cycles i7 index, 10 cycles i5 index, 90 cycles read2) at a targeted depth of 20,000 reads per nucleus.

###### Single nuclei sequencing analysis

Preprocessing of sequencing data was performed with Cell Ranger^99^ version 7.0.1, aligning to the GRCm38 mouse reference genome with the ‘--include-introns’ flag. We excluded cells with a low number of detected genes (ngenes), keeping cells with ngenes > 200 per sample, as established in previous studies^100^.

After filtering out lowly-expressed genes, we merged all sample datasets. We selected the top 7000 most differentially expressed genes across samples to be analyzed in the dataset, then proceeded with log2normalization, centering, principal component analysis with 100 components (PCA), and Uniform Manifold Approximation and Projection (UMAP). Data was clustered using the Louvain method^101^ and annotated using canonical markers of cellular identity^102^. Two-dimensional embeddings for visualization were created using UMAP.

To analyze microglial subpopulations, microglial data was subset from original, pre-processed objects, and re-normalized/centered/PCA/and UMAP-analyzed as described. Microglial subpopulations were clustered using the Louvain method and annotated based on established microglial markers^25, 26, 103^. Microglial gene signature scores were calculated incorporating these previously-established microglial signatures and averaging the expression of overlapping genes in our dataset. P value significance for microglial differential gene expression analysis was determined following the edgeR package^104^ to calculate log2foldchange (Log2FC) with multiple testing correction (Benjamini-Hochberg False Discovery Rate, “FDR”). Genes with FDR < 0.05 are considered significantly differentially expressed. Microglial subpopulation abundance was calculated as the number of each microglial subtype out of the total microglia identified per sample, averaged across each experimental group and expressed as a percentage.

### QUANTIFICATION AND STATISTICAL ANALYSIS

For all quantifications, experimenters were blinded to sample identity and condition.

For fluorescent immunohistochemistry and confocal microscopy, images were taken at 20X or 63X and cells considered co-labeled when markers co-localized within the same plane. For microglia staining at least three sections per mouse, 3-5 frames in standardized locations per section were counted. In all experiments, “n” refers to the number of mice. In every experiment, “n” equals at least 3 mice per group; the exact “n” for each experiment is defined in the figure legends.

All statistics were performed using Prism Software (Graphpad). For all analyses involving comparisons of one or two variables, 1-way or 2-way ANOVAs, respectively, were used with Tukey’s multiple comparisons post hoc corrections used to assess main group differences. For analyses involving only two groups, unpaired two-tailed Student’s t tests were used. Shapiro-Wilk test was used to determine normality for all datasets; all datasets were parametric. A level of p < 0.05 was used to designate significant differences.

---

**Supplementary Figure 1 - related to Figure 3**

**Supplementary Figure 1:**
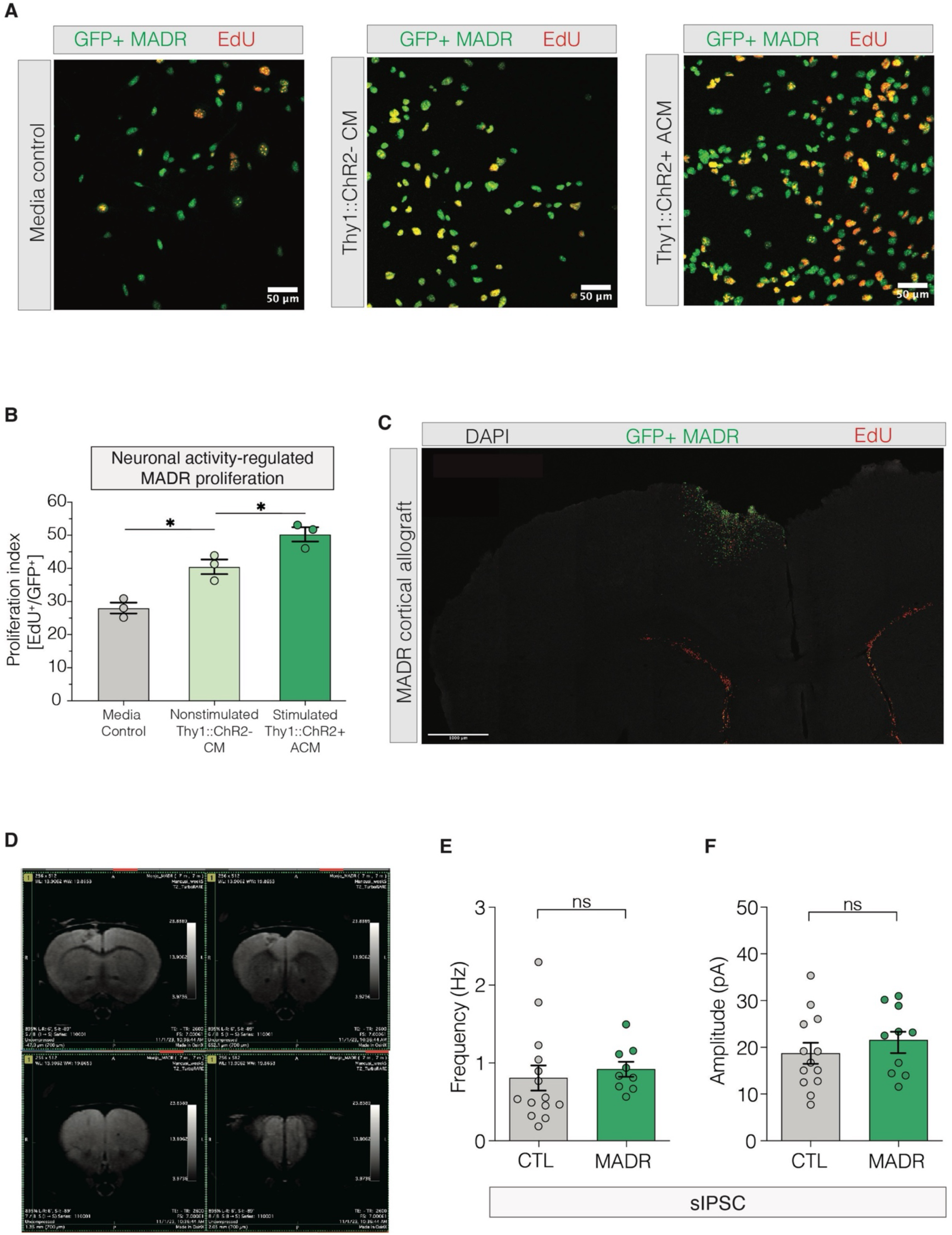
Murine glioma model, MADR, conserves human glioma biological and electrophysiological features. (A) GFP+ MADR cells (green) were incubated with aCSF (“Media control”) or ex vivo generated conditioned media from nonstimulated slices (“Thy1::ChR2-”) or from stimulated slices (“Thy1::ChR2+”). EdU (5-ethynyl-2’-deoxyuridine, red) marks dividing cells. Scale bar, 20 um. (B) MADR cell proliferation was measured as the % of EdU+ cells of the total GFP+ MADR cells (“Proliferation Index”, y-axis), following exposure to aCSF (“Media control”), ex vivo generated conditioned media from nonstimulated slices (“Thy1::ChR2-”), or from stimulated slices (“Thy1::ChR2+”). (C) Representative histology of cortical allografts with GFP+ MADR cells (green). DAPI (4’,6-diamidino-2-phenylindole, blue) marks nuclei, EdU (red) marks dividing cells. Scale bar, 1000 um. (D) Representative T2-enhanced MRI images (magnetic resonance imaging) of cortical MADR allografts. (E-F) Frequency and amplitude of spontaneous inhibitory postsynaptic currents (sIPSCs) were measured. Each individual point represents the average proliferation index of a single culture of MADR cells (B). Data are presented as mean ± s.e.m. One-way ANOVA with Tukey’s post hoc analysis (B) or unpaired t test with Welch’s correction (E-F); *P < 0.05; NS, not significant.

**Supplementary Figure 2 - related to Figure 3**

**Supplementary Figure 2:**
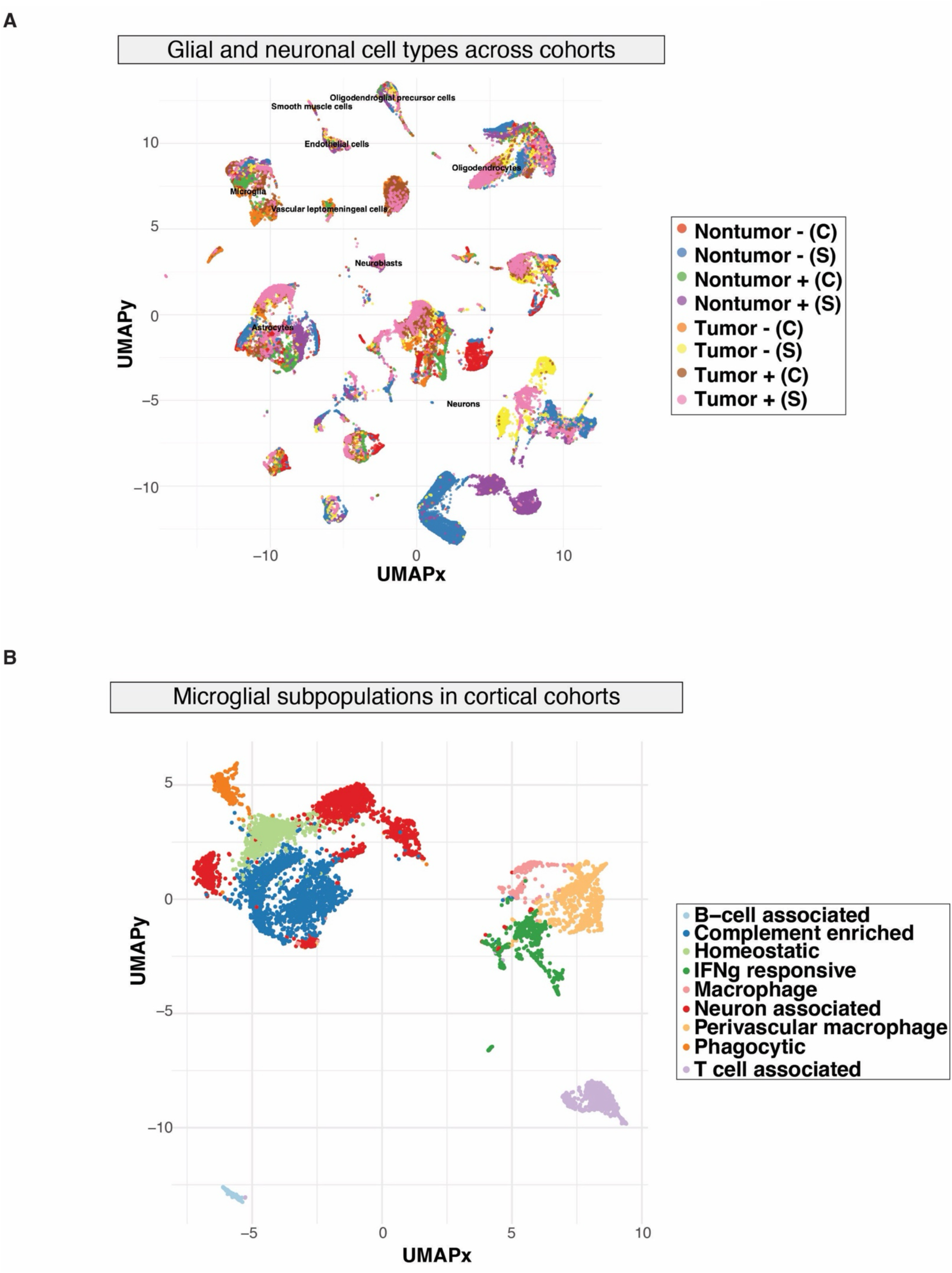
Multiple cellular subpopulations identified in single nuclei sequencing dataset. (A) Single nuclei sequencing was performed on tissue obtained from glioma allografted (“Tumor”) or saline-injected (“Nontumor”) control mice, which received cortical optogenetic stimulation (Thy1::ChR2+, “+”) or nonstimulated blue light control (Thy1::ChR2-, “-”). Both cortical (“C”) and striatal (“S”) tissue was collected from all cohorts of animals. Following Seurat-based computational analysis, single cells were mapped in a dimensionality-reduced context in 2 reduced dimensions, “UMAPy” (y-axis) and “UMAPx” (x-axis). (B) Cells belonging to the microglial cellular cluster comprise multiple subpopulations of microglial states, with single microglial cells mapped in 2 reduced dimensions, “UMAPy” (y-axis) and “UMAPx” (x-axis). Each point represents a single cell captured in the dataset, clustering in unique cellular populations (labeled on plot, A-B).

**Supplementary Figure 3 - related to Figure 4**

**Supplementary Figure 3:**
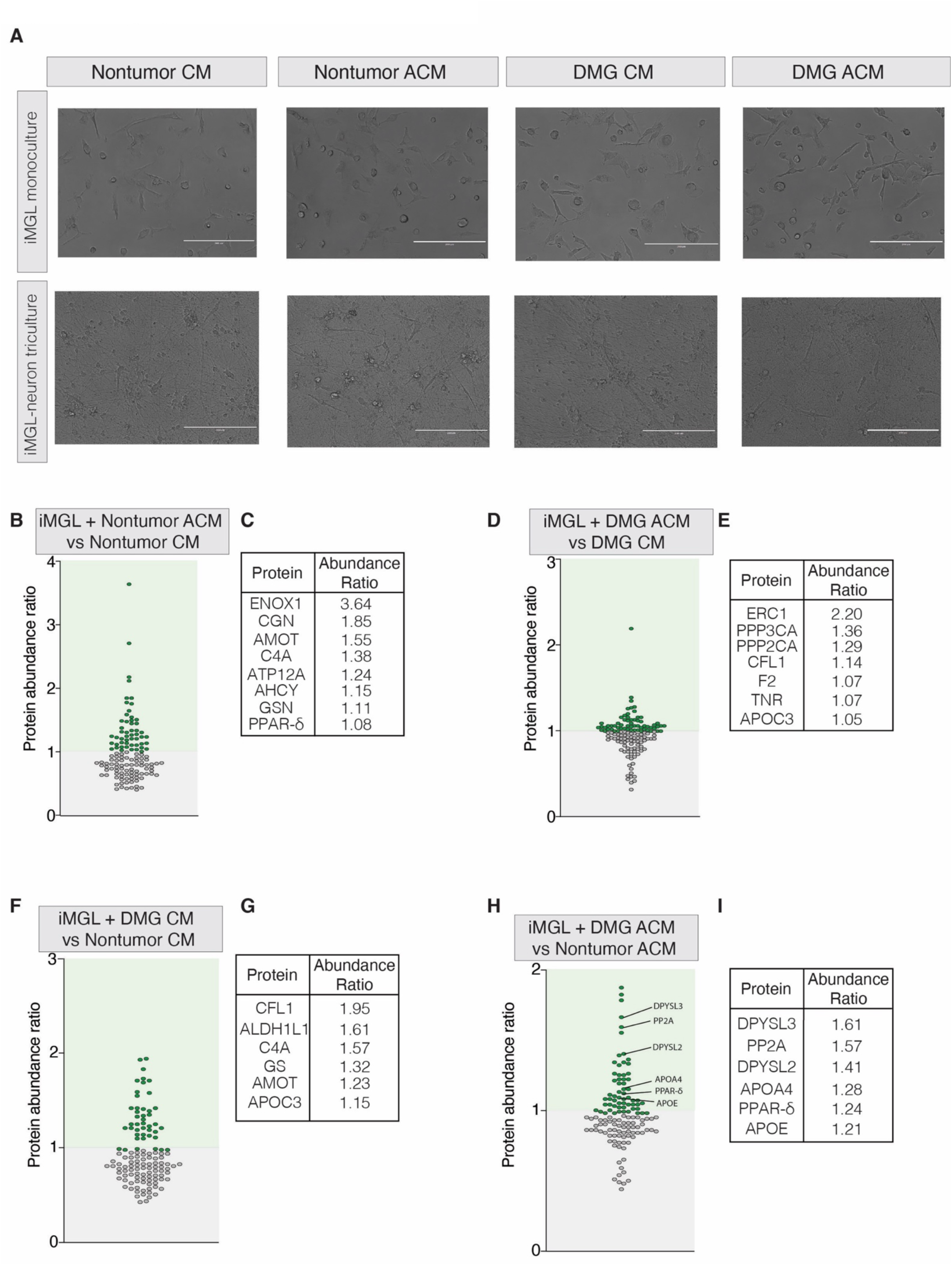
Human microglia respond to glioma-secreted factors. (A) Human iPSC-derived microglia (“iMGL”) in both monoculture and neuron tri-culture change morphology in response to acute exposure to ex vivo generated conditioned media from nontumor, nonstimulated (“Nontumor CM”), nontumor, stimulated (“Nontumor ACM”), glioma-bearing nonstimulated (“DMG CM”), or glioma-bearing stimulated (“DMG ACM”) slices. Scale bar, 200 um. (B-I) Supernatant from iMGL monocultures underwent LC/MS analysis following acute exposure to the aforementioned ex vivo conditioned media, calculating the ratio of protein abundance (“Protein Abundance Ratio”, y-axis) normalized on total peptide amount. Ratios are calculated by comparing iMGL supernatant after exposure to nontumor ACM and nontumor CM (B-C), glioma ACM and glioma CM (D-E), glioma CM and nontumor CM (F-G), and glioma ACM and nontumor ACM (H-I).

**Supplementary Figure 4 - related to Figure 4**

**Supplementary Figure 4:**
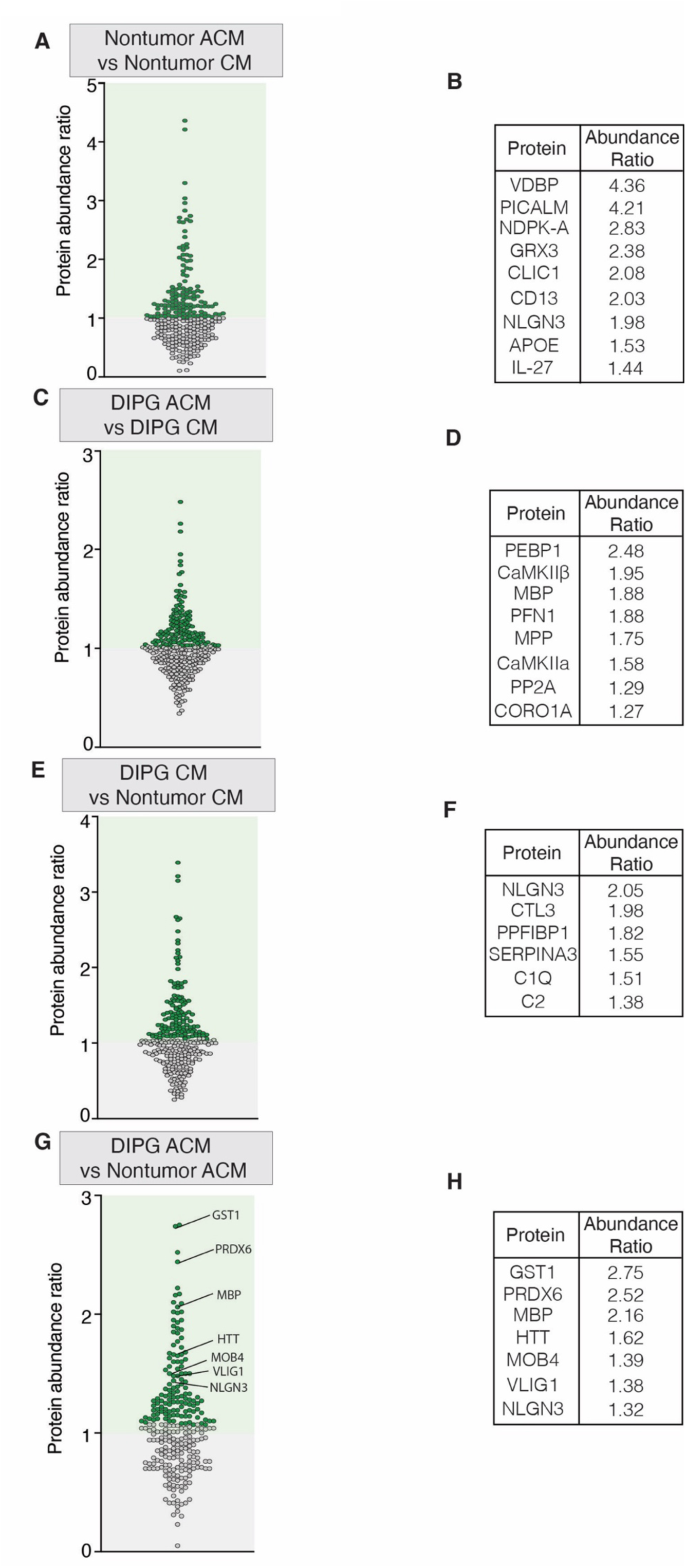
Comprehensive analysis of ex vivo slice generated secretome. (A-H) Ex vivo slice generated conditioned media underwent LC/MS analysis, calculating the ratio of protein abundance (“Protein Abundance Ratio”, y-axis) normalized on total peptide amount. Ratios are calculated by comparing ex vivo nontumor ACM and nontumor CM (A-B), glioma ACM and glioma CM (C-D), glioma CM and nontumor CM (E-F), and glioma ACM and nontumor ACM (G-H).

**Supplementary Table 1**: Microglial subpopulation gene enrichment list.

The top 50 most significantly-upregulated (p<0.01) genes associated with each microglial signature (B-cell associated, complement-enriched, homeostatic, IFNg responsive, macrophage, neuron associated, perivascular macrophage, phagocytic, and T cell associated, as in Fig. 3) is listed.

**Supplementary Video 1**: Microglial engulfment of postsynaptic structures in cortical xenograft model of pediatric high-grade glioma.

HNA+ glioma cells (SU-DIPG-VI, violet), Iba1+ microglia (white), and mCherry+ gephyrin+ synapses (red) in xenografted deep-layer premotor cortex of Thy1::ChR2 negative (“ChR2-”) mice. Scale bar, 10 um.

**Supplementary Video 2**: Human microglia engulf neuron-neuron postsynaptic structures in tri-culture with human GABAergic and glutamatergic neurons, following exposure to DMG-secreted factors.

Microglia (white), neurons (blue), PSD95+ excitatory synapses (red), and gephyrin+ inhibitory synapses (green) following treatment with conditioned media generated from stimulated glioma-bearing slices (“DMG ACM”). Scale bar, 20 um.

